# Sequential CRISPR gene editing in human iPSCs charts the clonal evolution of leukemia

**DOI:** 10.1101/2020.04.21.051961

**Authors:** Tiansu Wang, Allison R Pine, Josephine Wesely, Han Yuan, Lee Zamparo, Andriana G Kotini, Christina Leslie, Eirini P Papapetrou

## Abstract

Human cancers arise through an evolutionary process whereby cells acquire somatic mutations that drive them to outgrow normal cells and create successive clonal populations. “Bottom-up” human cancer evolution models could help illuminate this process, but their creation has faced significant challenges. Here we combined human induced pluripotent stem cell (iPSC) and CRISPR/Cas9 technologies to develop a model of the clonal evolution of acute myeloid leukemia (AML). Through the sequential introduction of 3 disease-causing mutations (ASXL1 C-terminus truncation, SRSF2^P95L^ and NRAS^G12D^), we obtained single, double and triple mutant iPSC lines that, upon hematopoietic differentiation, exhibit progressive dysplasia with increasing number of mutations, capturing distinct premalignant stages, including clonal hematopoiesis, myelodysplastic syndrome, and culminating in a transplantable leukemia. iPSC-derived clonal hematopoietic stem/progenitor cells recapitulate transcriptional and chromatin accessibility signatures of normal and malignant hematopoiesis found in primary human cells. By mapping dynamic changes in transcriptomes and chromatin landscapes, we characterize transcriptional programs driving specific stage transitions and identify vulnerabilities for early therapeutic targeting. Such synthetic “de novo oncogenesis” models can empower the investigation of multiple facets of the malignant transformation of human cells.

## Introduction

Cancer develops through the successive accumulation of somatic mutations that drive a reiterative process of clonal selection and evolution. It has long been recognized that understanding this process could guide early intervention, but opportunities to comprehensively study it have been limited^1^. Human cancers typically take years to develop and require multiple hits, in stark contrast to mouse cells that transform much faster with only one or few mutations. Human models of “de novo oncogenesis” that recapitulate the conversion of a normal cell into a malignant cell in a stepwise manner could provide important insights, but their development has faced major challenges: the limited ability of primary human cells for ex vivo growth to allow sufficient time for natural or artificial selection of clonal populations harboring oncogenic mutations; their inefficient gene modification; and gaps in knowledge on the sets of mutated genes that are necessary and sufficient to transform specific human cell types. Previous attempts at transforming human cells in the lab employed strong viral oncogenes and/or ectopic expression of cellular oncogenes^2–4^. More recently, CRISPR-mediated mutagenesis in human intestinal organoids was used to model colorectal cancer^5,6^.

Similar to most cancers, acute myeloid leukemia (AML) results from the sequential acquisition of driver mutations by the same hematopoietic stem/progenitor cell (HSPC) clone over time. Large-scale sequencing of tumor genomes of patients with AML, as well as individuals with clinically defined preleukemic conditions – myelodysplastic syndrome (MDS) and age-related clonal hematopoiesis (CH) – have yielded crucial information on the relative timing of acquisition of specific gene mutations^7,8^. More recently, sequencing of large AML patient cohorts revealed robust mutational cooperation patterns enabling inference of mutational paths^9^. However, even in cancers with clinically recognizable intermediate states and accessible tumor material, like AML, clonal heterogeneity and genetic background differences among patients severely limit the types of analyses that can be performed with primary tumor cells.

Here we exploit precise CRISPR-mediated gene editing and the amenability of human iPSCs to clonal selection to reconstruct the clonal evolution of AML. We show that HSPCs derived from isogenic iPSC lines exhibit progressive malignant features with increasing numbers of mutated genes, capturing CH, MDS and AML stages, with the latter endowing engraftment ability of immature human blasts into immunodeficient mice, and recapitulate primary AML in morphology, gene expression and chromatin accessibility. Leveraging the ability to obtain homogeneous populations of clonal iPSC-HSPCs upon hematopoietic differentiation, we perform integrative genomics analyses of their transcriptome and chromatin landscapes and identify AML targets for early intervention.

## Results

### Gene edited iPSCs phenotypically capture the clonal evolution of AML

We selected a prototypical secondary AML (sAML, i.e. arising from preexisting MDS), corresponding to the “chromatin-spliceosome” mutational path^9^. This genomic category constitutes the most common genetic group of sAML and is characterized by a combination of mutations in genes encoding a chromatin regulator and a regulator of RNA splicing, together with late “signaling” mutations^9^. From within this group, we selected the most common combination of specific gene mutations based on patterns of cooccurrence from population genetics studies^9,10^. Specifically, we sequentially introduced the mutations: ASXL1 C-terminus truncation; SRSF2^P95L^; NRAS^G12D^, all in heterozygous state, using CRISPR-mediated gene editing of a normal human iPSC line (N-2.12, previously described and extensively characterized)^11^ (Fig. 1a and Extended Data Fig. 1a,b). Additionally, in order to model mutational redundancy, we further introduced FLT3-ITD as a fourth mutation, as RAS and FLT3 mutations are mutually exclusive in human AML^9^ (Extended Data Fig. 1c). Three independent clones from each step (P: parental WT; A: ASXL1-mutant; SA: ASXL1- and SRSF2- double mutant; SAR: ASXL1-, SRSF2- and NRAS- triple mutant; and SARF: ASXL1-, SRSF2-, NRAS- and FLT3- quadruple mutant) – generated by two different gRNAs to exclude the possibility of off-target events confounding cellular and molecular phenotypes – were isolated and analyzed after hematopoietic differentiation (Supplementary Table 1). These experiments revealed progressive phenotypic changes. A gradual loss of differentiation potential was observed from A to SAR, as demonstrated by retention of CD34 expression (Fig. 1b and Extended Data Fig. 2a), morphological features of immature myeloid progenitor cells (Fig. 1c) and decreased colony-forming ability (Fig. 1d). A and SA iPSC-HSPCs showed reduced growth in vitro, while growth rate was restored in SAR iPSC-HSPCs to normal or above normal levels (Fig. 1e). SAR and SARF iPSC-HSPCs had higher fractions of cells in S phase (Extended Data Fig. 2b,c) and markedly extended in vitro survival (over 70 days, while normal iPSC-HSPCs arrest proliferation after 21 days) (Fig. 1f and Extended Data Fig. 2d). The growth disadvantage of A and SA cells is consistent with our previous observations in MDS patient-derived iPSCs and previously reported findings in several models of SF mutations^11–14^.

**Fig. 1.**
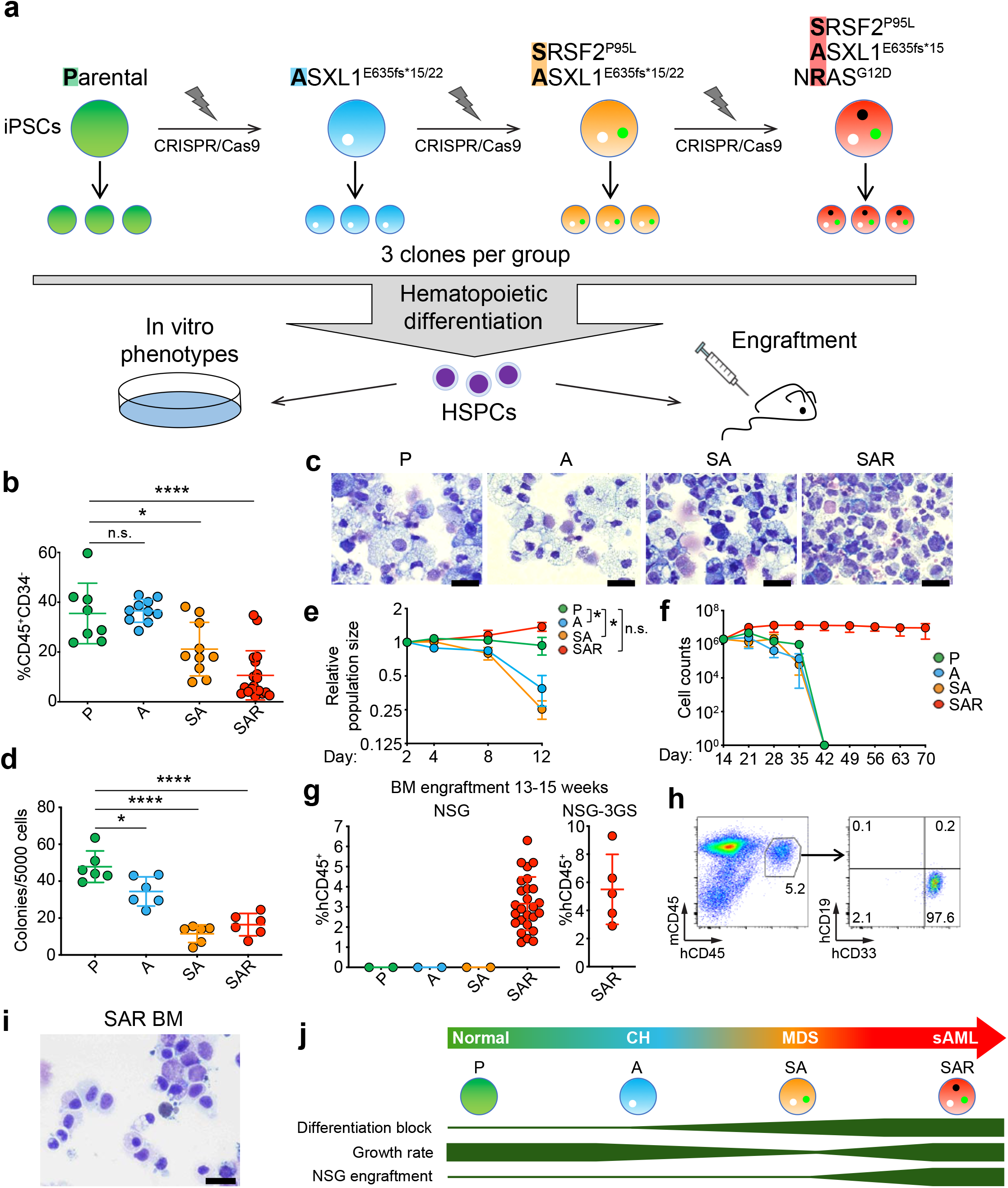
HSPCs generated from gene-edited iPSCs phenotypically mimic the clonal evolution of AML. **a,** Schematic overview of the generation and phenotypic characterization in vitro and in vivo of isogenic clonal iPSC lines generated through CRISPR/Cas9-mediated gene editing. Stepwise introduction of mutations was performed to generate isogenic single, double, triple and quadruple mutant iPSCs. Three independent lines per genotype were established after detailed genetic characterization (see also Supplementary Table 1, Extended Data Fig. 1 and Materials and Methods for details) and subjected to hematopoietic differentiation for phenotypic characterization and engraftment assays in NSG and NSG-3GS mice. **b,** Fraction of CD34^-^/CD45^+^ cells, i.e. hematopoietic cells that have lost CD34 expression upon maturation, on day 14 of hematopoietic differentiation. Mean and SEM of values from 5-12 independent differentiation experiments with 2 different iPSC lines from each genotype (A-1, A-2, SA-1, SA-2, SAR-1, SAR-2) are shown (see also Extended Data Fig. 2a). **c,** Wright-Giemsa staining of representative cytospin preparations of hematopoietic cells derived from the indicated iPSC lines after 14 days of hematopoietic differentiation culture. P, A and SA panels show mainly differentiated myeloid cells (granulocytes and monocytes), whereas the SAR panel shows immature myeloid progenitors with blast morphology. Scale bars, 25 μm. **d,** Number of colonies obtained from 5,000 cells seeded in methylcellulose assays on day 14 of hematopoietic differentiation. Mean and SEM of 2-4 independent methylcellulose experiments using 2 different iPSC lines from each genotype are shown (A-1, A-2, SA-1, SA-2, SAR-1, SAR-2). **e,** Competitive growth assay. The cells were mixed 1:1 at the onset of hematopoietic differentiation with an iPSC line stably expressing GFP derived from the parental line. On days 2-12 of differentiation the relative population size was estimated as the percentage of GFP^-^ cells (calculated by flow cytometry) at each time point relative to the population size on day 2. Mean and SEM of 2-3 independent experiments using 1 or 2 different iPSC lines from each genotype are shown. **f,** Cell counts of HSPCs at the indicated days of liquid hematopoietic differentiation culture. Mean and SEM of 2-3 independent differentiation experiments with 1 or 2 different iPSC lines from each genotype are shown. **g,** Levels of human engraftment in the BM of NSG and NSG-3GS mice 13 to15 weeks after transplantation with HSPCs derived from one iPSC line per genotype (P: N-2.12, A: A-1, SA: SA-2, SAR: SAR-1). Error bars show the mean and SEM of individual mice. For SAR, values from 26 NSG and 5 NSG-3GS mice from 5 independent transplantation experiments are included. **h,** Representative flow cytometry panels assessing human cell engraftment and lineage markers in the BM of recipient mice 13-15 weeks posttransplantation. **i,** Wright-Giemsa-stained cytospin of BM cells of a recipient mouse transplanted with SAR hematopoietic cells. Scale bar, 25 μm. **j,** Schematic summary of all in vitro and in vivo phenotypic analyses. (P: Parental, A: ASXL1^C-truncation^, SA: SRSF2^P95L^ /ASXL1^C-truncation^, SAR: SRSF2^P95L^ /ASXL1^C-truncation^/ NRAS^G12D^)

We transplanted iPSC-HSPCs derived from the gene edited cells into NOD/SCID-IL-2Receptor-γchain-null (NSG) and NSG-3GS (engineered to constitutively produce human IL-3, GM-CSF and SCF^15^) mice. Normal (P), A and SA iPSC-HSPCs showed no detectable engraftment. In contrast, all 31 mice transplanted with SAR cells had detectable engraftment of human hematopoietic cells at levels ranging from 1.2% to 6.3% in NSG and up to 9.3% in NSG-3GS mice 13-15 weeks post-transplant and enlarged spleens (Fig. 1g and Extended Data Fig. 2e,f). All human cells engrafted were of myeloid lineage (CD33+) and blast morphology (Fig. 1h,i), consistent with leukemic engraftment.

No significant phenotypic differences between SAR and SARF iPSC-HSPCs were observed in any of the in vitro or in vivo assays (Extended Data Fig. 2), providing no evidence of enhancement of the leukemic phenotype of SAR cells by the FLT3-ITD mutation, as expected. These results support redundancy of constitutive FLT3 signaling with RAS pathway activation in driving AML and are consistent with the lack of co-occurrence of these “signaling” mutations in the same clone in AML patients^9^.

Collectively, these in vitro and in vivo phenotypes recapitulate malignant features similar to those that we previously characterized in patient-derived iPSC lines capturing distinct disease stages from CH to MDS and AML^12,16^, with single mutant cells representing a CH stage, double mutant cells an MDS stage and triple mutant cells a leukemia (Fig. 1j).

### Gene edited iPSCs capture transcriptional and chromatin landscapes of primary AML

We next isolated CD34+/CD45+ iPSC-HSPCs from 2-3 clones of each group and performed RNA-seq and ATAC-seq analyses (Extended Data Fig. 3a,b). Principle component analysis (PCA) of both transcriptome and chromatin accessibility data grouped the samples by genotype with principle component 1 (PC1) capturing roughly disease progression both in RNA-seq and ATAC-seq datasets (Fig. 2a,b). SAR and SARF grouped closer to each other than other genotypic groups in both analyses, consistent with their close phenotypic similarities. Gene set enrichment analysis (GSEA) showed enrichment of gene expression signatures of primary AML, and of “chromatin-spliceosome” AML subgroup more specifically, in SAR and SARF cells (Extended Data Fig. 4a,b). Furthermore, several genes found in single-cell transcriptome analyses of primary cells to be preferentially expressed in AML HSPC-like cells relative to normal HSPCs from healthy bone marrow (BM), were found upregulated in SAR and SARF HSPCs (Fig. 2c)^17^. Conversely, genes preferentially expressed in primary normal differentiated cells of the myeloid lineage were found downregulated in SAR and SARF compared to P, A and SA cells (Fig. 2d). GSEA using a ranked list of genes linked to chromatin regions more accessible in SAR cells also revealed enrichment for early hematopoietic progenitor gene sets (Extended Data Fig. 4c). We also compared the chromatin accessibility landscapes of our cells to those defined in primary human hematopoietic cell types along the hematopoietic hierarchy^18^. The chromatin landscapes of P, A and SA cells predominantly resembled those of granulocyte-monocyte progenitors (GMP), whereas SAR and SARF cells more closely matched earlier progenitors (common myeloid progenitors, CMP and multipotent progenitors, MPP) and HSCs (Fig. 2e). Furthermore, we found peaks associated with AML HSPC genes^17^ to be more accessible in SAR and SARF HSPCs (Extended Data Fig. 4d).

**Fig. 2.**
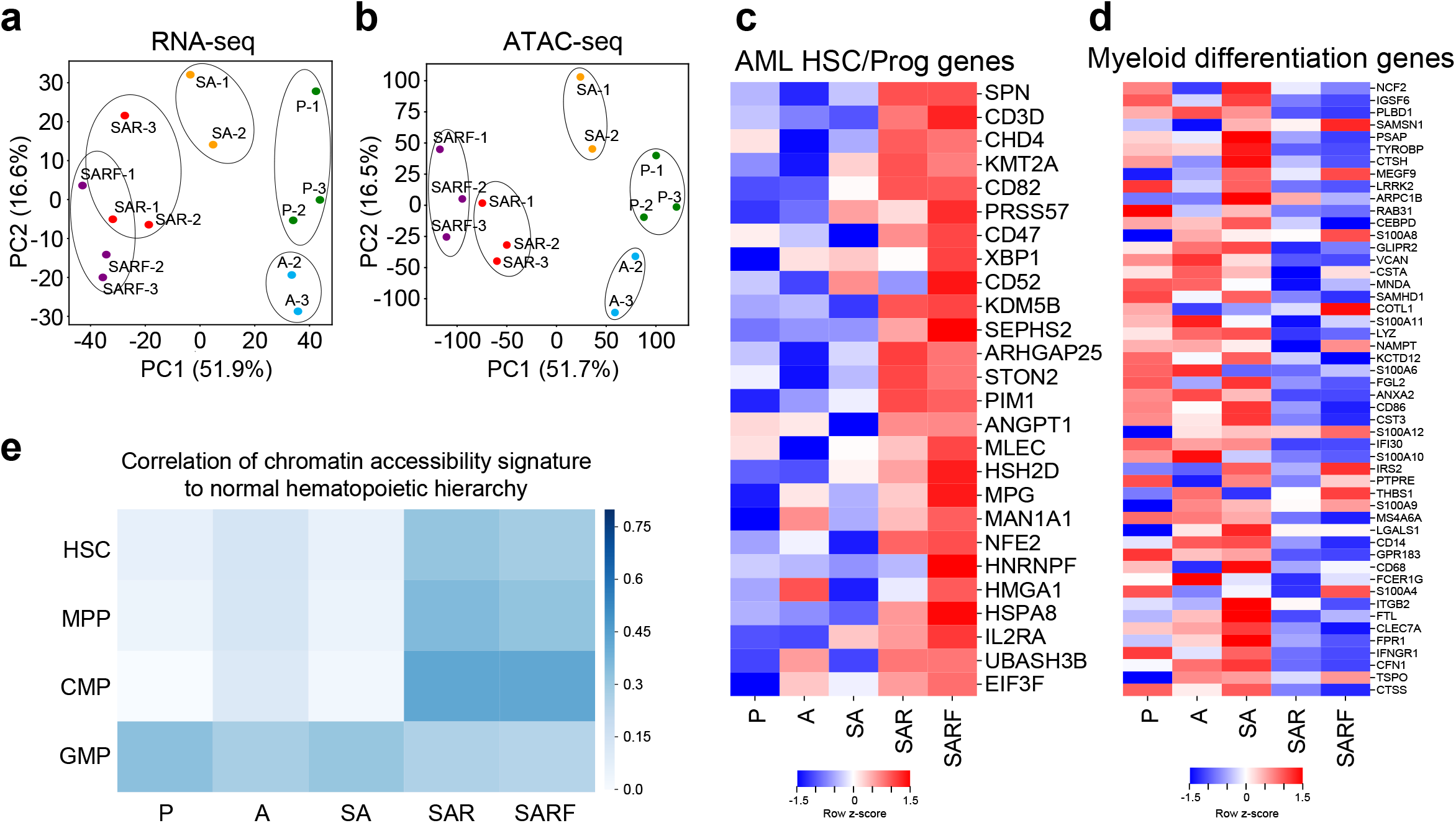
Transcriptional and chromatin accessibility landscapes of gene-edited iPSC-HSPCs capture those of primary human cells and show leukemic features. **a,** Principalcomponent analysis (PCA) of RNA-seq data. **b,** PCA of ATAC-seq data. **c,** Heatmap showing expression of genes preferentially expressed in primary AML HSPCs relative to normal HSPCs from healthy BM^17^, showing marked upregulation in SAR and SARF stages. **d,** Heatmap showing expression of a set of myeloid differentiation genes derived from normal primary cells^17^, showing marked downregulation in SAR and SARF. **e,** Heatmap showing Pearson correlation values of normalized read counts for ATAC-seq peaks that overlap between our clonal evolution stages (P, A, SA, SAR, SARF) and primary normal hematopoietic cell subpopulations (hematopoietic stem cell, HSC; multi-potent progenitor, MPP; common myeloid progenitor, CMP; granulocyte-monocyte progenitor, GMP) from Corces et al.^18^.

Collectively, these data show that gene edited iPSC-HSPCs recapitulate gene expression and chromatin accessibility changes found in primary human normal hematopoietic cells along the differentiation hierarchy and in primary AML cells, with SAR and SARF HSPCs resembling immature progenitors with transcriptional and chromatin features of AML.

### Dynamics of transcriptional and chromatin accessibility changes during clonal evolution

We next took advantage of our model to characterize dynamic changes of gene expression and chromatin accessibility along the process of clonal evolution. Changes in gene expression and chromatin accessibility in both directions were observed in all stages, with changes accumulating at each progressive stage (Fig. 3a,b, Extended Data Fig. 4e-g). To globally assess the directionality and persistence of changes over the course of disease progression, we plotted all gene expression and all chromatin accessibility changes (log2FC) at each stage (relative to P) against those in the subsequent stages (Fig. 3c,d and Extended Data Fig. 4h,i). The majority of gene expression and chromatin accessibility changes found in the A stage (compared to P) did not persist in SA nor in the subsequent stages (Fig. 3c,d and Extended Data Fig. 4h,i). In contrast, gene expression and chromatin accessibility changes were more concordant between the SA and SAR stages and – even more so – between the SAR and SARF stages (Fig. 3c,d). This analysis indicates that early (A stage) changes are mostly transient, while a higher proportion of the changes established at the SA stage and thereafter persist in the subsequent stages.

**Fig. 3.**
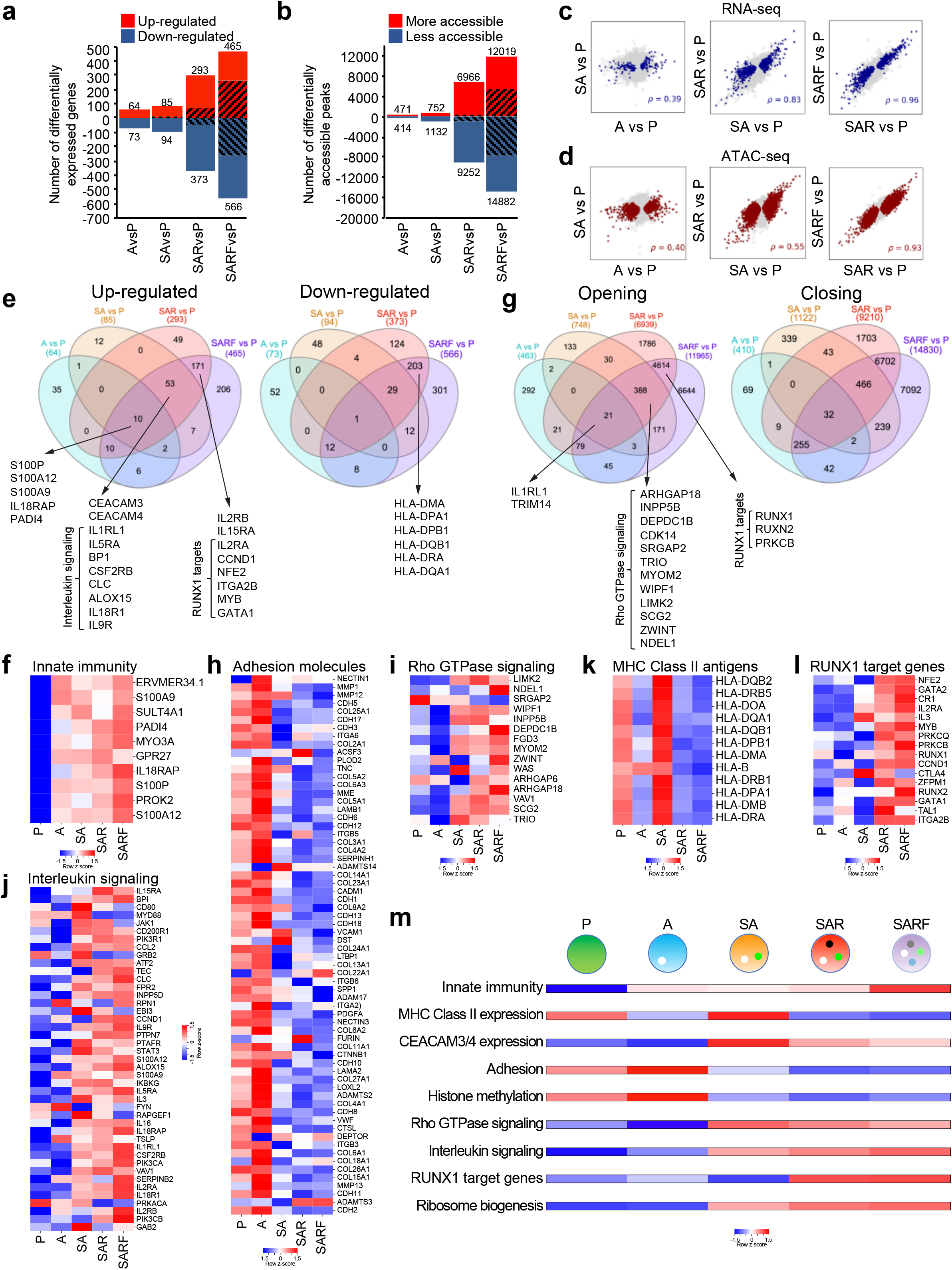
Dynamic gene expression and chromatin accessibility changes during clonal evolution. **a,** Bar plot showing the number of differentially expressed genes (DEGs, log2FC > 1, FDR-adjusted p < 0.05) in the indicated comparisons. Shaded areas represent DEGs found to also be differentially expressed in a previous stage. **b,** Bar plot showing the number of differentially accessible peaks (DAPs, FDR-adjusted p < 0.05) in the indicated comparisons. Shaded areas represent DAPs found to also be differentially accessible in a previous stage. **c,** Scatter plots of gene expression changes (log2FC) showing correlation between the two comparisons indicated in the x and y axes of each plot. Significant changes (FDR-adjusted p < 0.05) in the x-axis comparisons are highlighted in blue. The ρ values represent Spearman correlation values calculated from the log2FC of genes with significant changes in the x-axis comparison. **d,** Scatter plots of chromatin accessibility changes (log2FC) showing correlation between the two comparisons indicated in the x and y axes of each plot. Peaks with a significant change (FDR-adjusted p < 0.05) in the x-axis comparisons are highlighted in red. The ρ values are Spearman correlation values calculated from the log2FC of peaks with significant changes in the x-axis comparison. **e,** Venn diagrams showing overlap of up- and down-regulated genes in the indicated comparisons with selected genes annotated. **f,** Heatmap showing the expression of the 10 upregulated genes that are common in all comparisons (A vs P, SA vs P, SAR vs P and SARF vs P, shown in the Venn diagram in E, left panel) across stages. Marked in red are genes involved in inflammatory responses and innate immunity. **g,** Venn diagrams showing overlap of opening or closing peaks in the indicated comparisons. Selected genes associated with the indicated groups of peaks are annotated. **h-l,** Heatmaps showing expression of the indicated gene sets (adhesion molecules, genes involved in Rho GTPase signaling, interleukin signaling, MHC Class II genes and RUNX1 target genes) across stages. **m,** Summary dynamics of transcriptional changes in selected gene sets, as indicated. The bar corresponding to each stage is colored by the median log-transformed gene expression level in each gene set.

Such persistent changes involved only 11 genes (10 up- and 1 down- regulated), which were significantly differentially expressed in A vs P and remained so in all subsequent stages (Fig. 3e,f). These included several genes involved in innate immunity, notably genes encoding members of the S100 family of calcium-binding proteins, including S100A9, whose role in early MDS pathogenesis has been proposed in several recent studies^19–21^ (Fig. 3f). Similarly, we found only a small number of persistent differentially accessible peaks, i.e. common to all groups (A, SA, SAR and SARF) and genes associated with them included IL1RL1 (IL-33 receptor) and TRIM14, also involved in IFN signaling (Fig. 3g).

Differential gene expression and accessibility analyses, together with Gene Ontology (GO) and GSEA analyses, revealed additional notable patterns, summarized in Fig. 3m. Expression of CEACAM 3 and 4 was found consistently increased, starting at the SA stage, but a number of other genes related to adhesion, integrin signaling and extracellular matrix composition were downregulated from the SA stage and onwards (Fig. 3h and Extended Data Figs. 5a,b and 6a). Many genes related to Rho GTPase signaling, a pathway with well-documented roles in the maintenance of HSCs, were both upregulated and more accessible starting at the SA stage (Fig. 3g,i and Extended Data Fig. 6b). A number of genes related to interleukin signaling were also upregulated in SA, SAR and SARF cells (Fig. 3e,j and Extended Data Fig. 5c). Expression of several MHC class II human leukocyte antigen (HLA) genes was downregulated in A, markedly upregulated in SA, and again downregulated in the SAR and SARF stages (Fig. 3e,k and Extended Data Fig. 6c). Downregulation of MHC class II molecules has been reported in patients with relapsed AML and may be an important mechanism of immune escape^22^. GSEA analyses also revealed negative regulation of histone methylation beginning at the SA stage (Extended Data Figs. 5d and 6d). Changes found at later stages (SAR and SARF) were the upregulation of several target genes of the RUNX1 transcription factor (TF), which were also associated with chromatin regions of increased accessibility in SAR and SARF cells (Fig. 3e,g,l and Extended Data Fig. 6e). Genes associated with increased ribosome biogenesis, including rRNA transcription, were also more expressed and more accessible in the SAR and SARF stages (Extended Data Fig. 5e,f). RUNX1 plays important roles in hematopoiesis and leukemia and has been shown to regulate ribosome biogenesis in HSPCs^23^.

### TF programs during clonal evolution

We next performed TF motif enrichment analyses, which revealed dynamic changes at different stages (Fig. 4a,b). Peaks with IRF motifs closed early at the A stage and remained closed throughout all subsequent stages (Fig. 4b). Peaks containing AP-1 motifs (JUN, FOS, BATF), as well as peaks containing NFkB motifs, gained accessibility during the P-to-A stage transition, but were subsequently closed and remained closed thereafter (Fig. 4b). AP-1 and NFkB motifs were enriched in adhesion and histone methylation genes (Extended Data Fig. 7a) that were downregulated and closed from the SA stage onwards (Fig. 3h,m and Extended Data Figs. 5a,b,d and 6a,d), suggesting potential involvement of these TFs in their regulation. CEBP were the predominant motifs in peaks opening at the SA stage (Fig. 4a,b). Both CEBP and NFkB motifs were enriched in peaks associated with MHC Class II genes (Extended Data Fig. 7a), whose regulation mirrored the accessibility changes of these two motifs (Figs. 3k,m, 4b and Extended Data Fig. 6c). GATA, E2F and MECOM were among the top motifs associated with peaks gaining accessibility at the SAR stage and were enriched in genes associated with Rho GTPase signaling (Extended Data Fig. 7a).

**Fig. 4.**
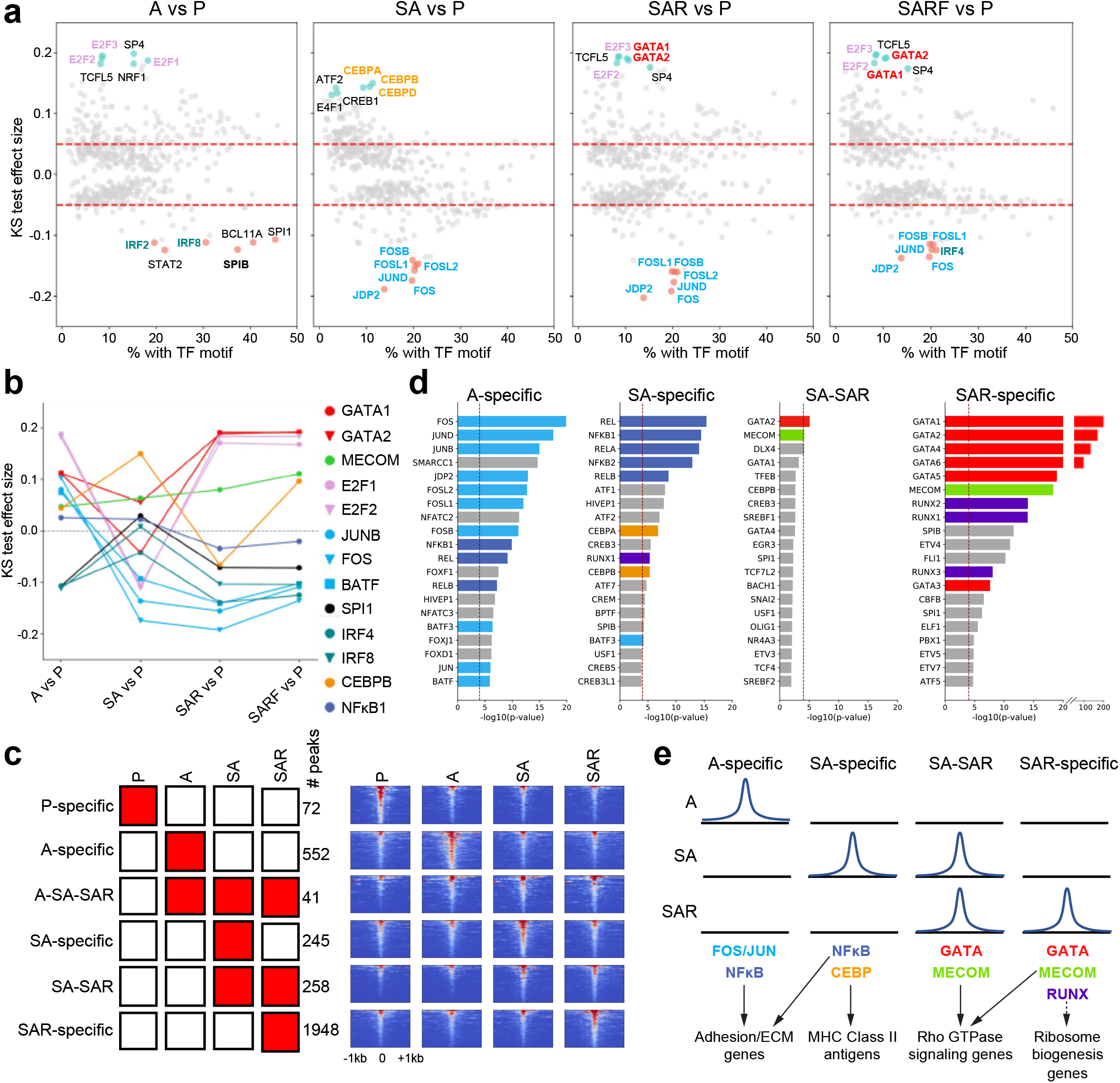
Characterization of transcriptional programs underlying distinct stage transitions. **a,** TF motif enrichment in the indicated stage transitions. Opening or closing peaks in each stage were compared with the total atlas and TF motif enrichments were calculated using the one-sided Kolmogorov-Smirnov (KS) test. The KS test effect size is shown on the y axis, and the percentage of peaks associated with the TF motif is plotted on the x axis. The dashed lines indicate TF motifs with a KS test effect size ≥ 0.05. The top 6 and bottom 6 motifs are marked in blue and red, respectively. **b,** TF motif enrichment dynamics across stages. Differentially accessible peaks in each stage compared to P were compared with the total atlas and TF motif enrichments were calculated using the one-sided KS test. The KS test effect size is shown on the y axis. **c,** Six stage-specific groups of ATAC-seq peaks defined based on their patterns of chromatin accessibility across the 4 stages P, A, SA and SAR. The number of peaks for each group is annotated. Left panel: schematic representation. Right panel: tornado plots of the annotated stage-specific groups of ATAC-seq peaks in the indicated stages. **d,** TF motif enrichment in the peaks of the 4 indicated stage-specific groups from a, containing numbers of peaks sufficient for motif enrichment analysis. The hypergeometric test was used to compare the enrichment of proportions of TF motifs for a stage-specific group (foreground ratio) versus those for the total atlas (background ratio). The top 20 most highly enriched TFs are shown. The red dashed vertical line indicates a Bonferroni-corrected p value threshold of 0.05. AP-1 TF family motifs are marked in light blue, NFkB family motifs are in dark blue, CEBP family motifs in orange, RUNX family motifs in purple, GATA family motifs in red and MECOM motifs in green. **e,** Summary of TF motif enrichment for the 4 stage-specific groups indicated (containing numbers of peaks sufficient for motif enrichment analysis), based on the data shown in d. Genes inferred to be putative relevant targets of these TFs (based on the analyses presented in Figs. 3, 4 and Extended Data Figs. 5, 6, 7a-c) are shown underneath, connected with arrows to the putative TFs. (A dashed arrow connects RUNX to ribosome biogenesis genes, as this link is inferred from the literature rather than from our data, which could not document enrichment for RUNX motifs in this gene set.)

To better identify TF programs acting at different stages, we sought to characterize dynamic chromatic changes in more detail. Clustering of all peaks that were differentially accessible in at least one stage compared to the parental revealed 2 clusters with overall increasing and 2 clusters with overall decreasing accessibility (Extended Data Fig. 7b,c). To identify TFs that drive specific stage transitions, we grouped ATAC-seq peaks based on their patterns of accessibility across stages into stage-specific groups (Fig. 4c). These groups contained similar relative proportions of intronic, promoter, exonic, and intergenic regions (Extended Data Fig. 7d). Analyses of the association between specific TF motifs and stage-specific groups of chromatin peaks revealed association of specific stages and stage transitions with distinct TF motifs (Fig. 4d,e). FOS/JUN and NFkB motifs were again found associated with open sites during the early A stage. GATA and MECOM motifs were associated with peaks opening at the SA-to-SAR transition and at the SAR stage. RUNX motifs were enriched in late-opening peaks acquired at the SAR stage.

### Identification of early events during AML establishment

These analyses also showed that the vast majority of chromatin sites that gained accessibility at the A stage (sum of: 552 A-specific peaks; 3 A-SA peaks; and 41 A-SA-SAR peaks; total 596) did not remain accessible in later stages (A specific, 552 peaks), consistent with our earlier analyses (Figs. 4c, 3d and Extended Data Fig. 4i). Approximately half of the sites that first became accessible at the SA stage (SA-specific and SA-SAR, total 503 peaks) remained accessible in SAR (SA-SAR group, 258 peaks) (Fig. 4c). Strikingly, the majority of SAR-accessible peaks (A-SA-SAR, SA-SAR and SAR-specific, total 2247 peaks) were acquired at the SAR stage (SAR-specific group, 1948 peaks, 86.7% of total), with only a small fraction established at earlier A or SA stages (A-SA-SAR and SA-SAR groups) (Figs. 4c, 5a). We reasoned that chromatin regions belonging to the SA-SAR group (which gained accessibility at the SA stage and remained accessible throughout the SAR and SARF stages) should contain the earliest molecular events among all the events that are important for AML establishment and maintenance (all SAR-accessible). Indeed, selecting genes associated with SA-SAR stagespecific peaks enriched for genes that are already open and upregulated at the SA stage (Fig. 5b, compare to Extended Data Fig. 7e and Supplementary Table 2). We thus hypothesized that genes selected from within this SA-SAR group could represent dependencies of AML that can enable AML targeting of an earlier disease stage. To test this, we first selected from all the 153 genes associated with the 258 SA-SAR peaks those that were most significantly upregulated in both SA and SAR (padj <0.1). We thus obtained 13 “AML-early” genes” (Fig. 5c, Supplementary Table 3). For validation of this selection approach, of those 13 genes, we selected *ATP6V0A2* because a specific inhibitor (bafilomycin-A1) is available and because its expression was found elevated in peripheral blood (PB) and BM from patients with AML (both de novo and secondary, as well as in “chromatin-spliceosome” AML), compared to healthy individuals (Fig. 5d-f)^24^. *ATP6V0A2* encodes a subunit of the vacuolar ATPase (v-ATPase), a proton transporter with emerging roles in cancer and drug resistance^25,26^. Consistent with our hypothesis, while AraC, which is the backbone of AML induction chemotherapy, preferentially killed SAR over SA HSPCs, as would be predicted by the higher proliferation rate of the former (Fig. 1e and Extended Data Fig. 2b,c), V-ATPase inhibition by bafilomycin-A1 killed both SAR and SA iPSC-HPCs (Fig. 5g).

**Fig. 5.**
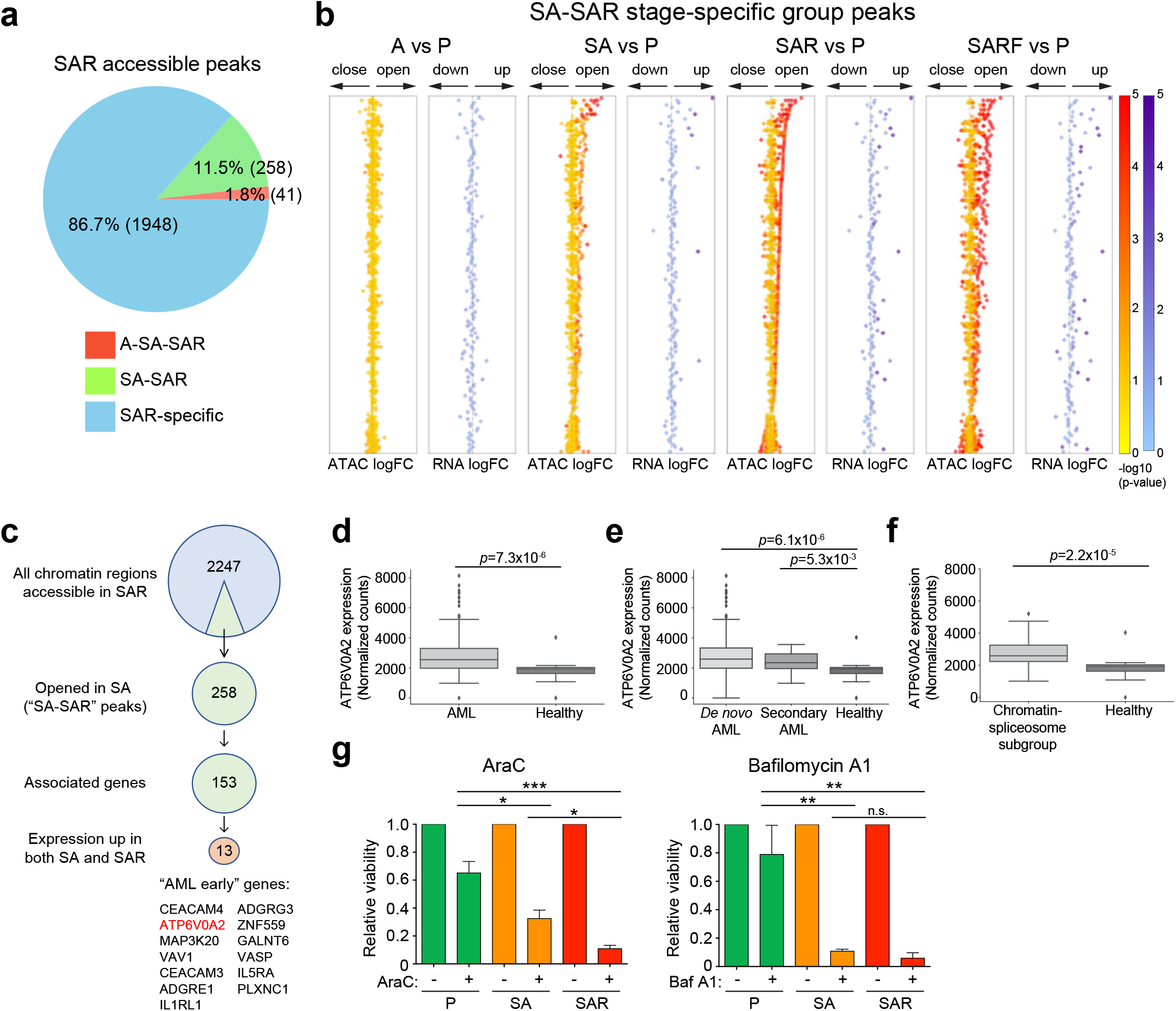
Identifying early AML targets. **a,** Breakdown of all peaks accessible at the SAR stage: A-SA-SAR peaks first become accessible at the A stage. SA-SAR peaks first become accessible at the SA stage. SAR-specific peaks only become accessible at the SAR stage. **b,** Diamond plots showing the accessibility change (log2FC) of the peaks belonging to the SA-SAR group together with the expression change (log2FC) of genes associated with them. Each row represents one gene and all peaks associated with it across all comparisons (A vs P, SA vs P, SAR vs P and SARF vs P), allowing back- and forward-tracking of changes in successive stages. **c,** Strategy for identification of “AML early” genes. From the 2247 ATAC-seq peaks that were accessible in SAR we selected the 258 that first became accessible at the SA stage and remained accessible in SAR (SA-SAR peaks). 153 genes could be associated with these 258 SA-SAR peaks. Of those, we selected 13 genes whose expression (RNA-Seq) was upregulated in both SA and SAR stages compared to P (padj < 0.1). The 13 genes are listed ranked by accessibility change in SAR vs P (See also Supplementary Table 3). **d-f,** Box plots of ATP6V0A2 expression (DESeq2-normalized counts) in PB and BM of patients from the BeatAML cohort^24^ grouped in de novo, secondary and “chromatin-spliceosome” AML, compared to that in healthy individuals. P values were calculated using a Wilcoxon rank-sum test. **g,** Left panel: Viability determined by cell counts of P, SA and SAR iPSC-HSPCs treated with AraC (200nM) compared to untreated cells. Right panel: Viability determined by cell counts of P, SA and SAR iPSC-HSPCs treated with the V-ATPase inhibitor bafilomycin A1 (Baf A1, 10nM) compared to untreated cells.

We further tested the preferential targeting of earlier clones through inhibition of *ATP6V0A2* over standard chemotherapy in iPSC lines directly derived from different clones of an AML patient. Cells from a patient with “chromatin-spliceosome” AML (AML-32) were reprogrammed and a clonal iPSC line representing a late AML clone (AML-32.13, harboring oncogenic mutations in *IDH2, ASXL1, SRSF2* and a “signaling” *CEBPA* mutation), as well as an iPSC line capturing an earlier clone in the evolution of AML (AML-32.18, harboring the *IDH2, ASXL1* and *SRSF2* mutations alone) were derived (Fig. 6a). AraC was more effective in killing the iPSC line AML-32.13 representing the later AML subclone, than the iPSC line AML-32.18 derived from the earlier AML clone, whereas bafilomycin-A1 had comparable effect in both clones (Fig. 6b). Finally, we tested V-ATPase inhibition, compared to AraC treatment, in primary “chromatin-spliceosome” AML cells from the same AML patient (AML-32) and from a second patient (AML-34) with *ASXL1, SRSF2, STAG2* clonal mutations and *CSF3R* subclonal mutation, using mutational variant allele frequencies (VAFs) to track the effects on different clones (Fig. 6c-g). V-ATPase inhibition resulted in a greater decrease in the SRSF2 mutation VAF than AraC, whereas it decreased CSF3R VAF at comparable or greater levels to AraC, indicating superior targeting of the earlier clone by bafilomycin-A1 (Fig. 6d,g).

**Fig. 6.**
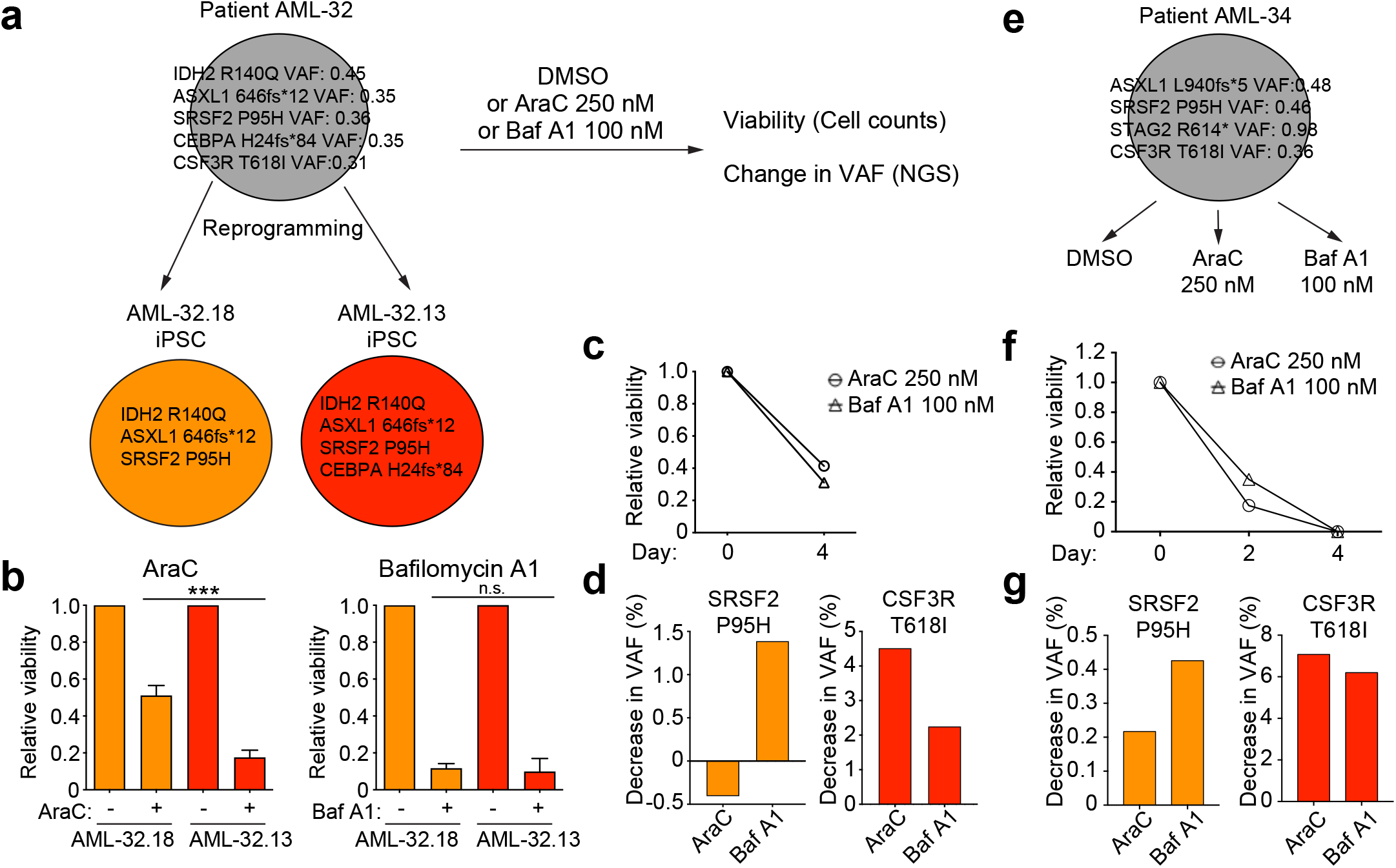
*ATP6V0A2* inhibition in primary AML cells. **a,** BM blasts from a “chromatin-spliceosome” AML patient (AML-32) were used to derive iPSC lines capturing both an early AML clone (AML-32.18, harboring only *IDH2, ASXL1* and *SRSF2* mutations) and a late AML clone (AML-32.13, harboring a *CEBPA* mutation, in addition to the *IDH2, ASXL1* and *SRSF2* mutations). **b,** Treatment of HSPCs derived from the iPSC lines shown in A with 200nM AraC (left) or 10nM bafilomycin A1 (right). AraC preferentially kills the later clone, while bafilomycin A1 has comparable efficacy in both clones. **c,** Viability of primary AML blasts from patient AML-32, treated with AraC or bafilomycin A1, determined by cell counts, relative to DMSO-treated cells. **d,** Changes in the clonal composition of the primary AML-32 patient cells after treatment with AraC or bafilomycin A1, assessed by measurement of the change in VAFs of selected mutations present in the sample. Left: decrease in the VAF of the clonal SRSF2 P95H mutation. Right: decrease in the VAF of the subclonal CSF3R T618I mutation. **e,** BM blasts from a “chromatin-spliceosome” AML patient (AML-34) were treated with AraC (250nM), bafilomycin A1 (100nM) or DMSO for 4 days. **f,** Viability of primary AML blasts from patient AML-34, treated with AraC or bafilomycin A1, determined by cell counts, relative to DMSO-treated cells. **g,** Decrease in the VAF of the clonal SRSF2 P95H mutation (left) and of the subclonal CSF3R T618I mutation (right) in primary AML blasts from patient AML-34 after treatment with AraC or bafilomycin A1, as indicated.

These data support the validity of our approach in identifying early AML targets that represent vulnerabilities common to both early and more evolved AML clones and that enable targeting of clones with truncal mutations more efficiently than standard chemotherapy.

## Discussion

Here we demonstrate the combination of CRISPR and human iPSC technology to build a human leukemia in a stepwise fashion, generating isogenic lines of increasing numbers of mutated genes and accordingly progressive malignant features. Our phenotypic findings are consistent with CH being associated with a single gene mutation in the vast majority of cases and MDS and AML typically requiring 2-3 and 3-5 mutations, respectively^9,10,27^.

Normal iPSC-HSPCs notoriously fail to engraft in xenograft assays^28^. We have previously shown that hematopoietic cells from iPSCs derived from patients with CH and MDS also fail to engraft^12^ and we therefore expected that P, A, and SA cells would not support engraftment, as our findings here indeed show. In striking contrast, we and others have shown that it is only iPSCs derived from AML patients that can produce HSPCs with engraftment potential^12,16^. This engraftment was predominantly myeloid, similarly to that of the SAR and SARF cells we present here, and unlike normal primary HSPC engraftment in the xenograft setting, which is heavily biased towards the B cell lineage^29^. Critically, AML-iPSCs are the only pluripotent stem cells (including embryonic stem cells and iPSCs) with demonstrated ability to engraft in xenografts upon in vitro hematopoietic differentiation to date. Thus, engraftment ability per se is the most characteristic feature of iPSC-derived leukemia described so far and we therefore used it here as the main surrogate AML phenotype.

Reassuringly, our phenotypic analyses here show that the gene edited iPSCs recapitulated the in vitro and in vivo phenotypes that we previously characterized in patient-derived iPSCs capturing all disease stages (encompassing disease-free, CH, low-risk MDS, high-risk MDS and sAML)^12^. Importantly, a critical limitation of the previous patient-derived progression models was the diverse genetic backgrounds of the different patients from which those iPSC lines were derived, which precluded analyses towards identifying stage-specific transcriptional and chromatin changes. In contrast, the isogenic conditions that we engineered here, enabled these comprehensive genomic analyses that were not possible with other models before.

Interestingly, the vast majority of both transcriptional and chromatin changes found at the single mutant stage (A) were transient. As ASXL1 encodes a chromatin regulator, these observations might reflect increased plasticity of the chromatin landscape caused by ASXL1 mutations, with most gene expression and chromatin accessibility changes constituting “epigenetic noise” (transient changes in our model) and only few required for the establishment of malignancy (persistent changes in our model)^30^. Of note, the few persistent changes involved genes implicated in innate immunity and inflammation. These findings may reflect observations of increased inflammatory responses that have recently been reported in patients with CH with *TET2* and *DNMT3A* mutations, especially since *ASXL1* is the third most commonly mutated CH gene after *TET2* and *DNMT3A*^27^. Importantly, while the increased inflammatory responses in the context of *TET2* and *DNMT3A* mutations that have previously been reported were found in mature immune cells, primarily monocytes, our results suggest that innate immunity and inflammatory signaling is already dysregulated at the HSPC level in CH. Interestingly, dysregulation of these pathways in cells early in the hematopoietic hierarchy have been suggested in the case of MDS^31,32^.

The main premise of studies into the clonal evolution of cancer is gaining insights that can inform targeting the disease early. This is particularly relevant for AML, where patient outcomes are mainly determined by relapse and resistance^33^. Most newly developed targeted therapies for AML (i.e. FLT3 inhibitors or IDH1/2 inhibitors) target mutations that are acquired late in the course of the disease. Because these are typically present in subclones and not in all leukemic cells of the patient, their targeting can eliminate subclones, but not the founding clone, which can give rise to residual disease or relapse. Our analyses reveal transcriptional programs that are critical for AML establishment and maintenance, with GATA2, MECOM and RUNX1 – TFs with previously recognized roles in AML – being the most prominent. Importantly, we present an analytical approach that takes advantage of the linearity of changes in our model to identify early events that are important to AML establishment (our “AML-early genes”) and provide proof of principle of the viability of this approach by targeting one of them (*ATP6V0A2*). We thus show that our iPSC de novo oncogenesis model can nominate AML vulnerabilities and provide a platform for functionally testing them in relevant cells.

While longitudinal studies of primary patient cells at different disease stages (CH, MDS, AML) have illuminated the mutational events underlying the clonal evolution of the disease, analyses of the type we present here are hindered by the clonal heterogeneity of primary material, particularly at the CH stage where mutant cells comprise just a small percentage (1-4%) of all cells. Mouse models of clonal evolution could conceivably be developed but an important caveat is that mouse cells typically require fewer events to transform and may thus not capture the multi-hit mutagenesis process underlying oncogenesis of human cells. In support of the relevance of our modeling approach to human AML, we show extensive similarities of gene expression and chromatin accessibility between our iPSC-derived cells and primary human hematopoietic cells, establishing that SAR and SARF iPSC-HSPCs resemble leukemic stem/progenitor cells. While some recent studies have provided evidence of non-linear (parallel or branching) clonal evolution in some cases of sAML^8,34^, a linear clonal evolution scenario remains the more likely mode of evolution in the majority of sAML cases, and is thus the most relevant and best-suited to our modeling approach.

In summary, by combining CRISPR and human iPSC technology, we built a human de novo leukemogenesis model that enabled us to chart the molecular events that underlie disease progression and identify molecular changes that are both critical for AML and occur early in the disease progression, and which can provide vulnerabilities for improved therapeutic targeting or biomarkers of disease progression.

## Acknowledgements

The authors thank Elli Papaemmanuil for advice on mutational selection. This work was supported by the US National Institutes of Health (NIH) grants R01HL137219 and R01CA225231, by the New York State Stem Cell Board, the Pershing Square Sohn Cancer Research Alliance and by a Scholar award from the Leukemia and Lymphoma Society. TW was supported by a Training Program in Stem Cell Biology fellowship from the New York State Department of Health (NYSTEM-C32561GG). AP was supported by the Tri-Institutional Training Program in Computational Biology and Medicine (CBM) funded by National Institutes of Health (NIH) grant 1T32GM083937.

## Author contributions

TW performed experiments and analyzed data. AP, HY, LZ and CL analyzed RNA-Seq and ATAC-seq data. AGK provided guidance. EPP conceived, designed and supervised the study, analyzed data and wrote the manuscript with assistance from TW and AGK.

## Supplementary Figure and Table legends

**Extended Data Fig. 1.**
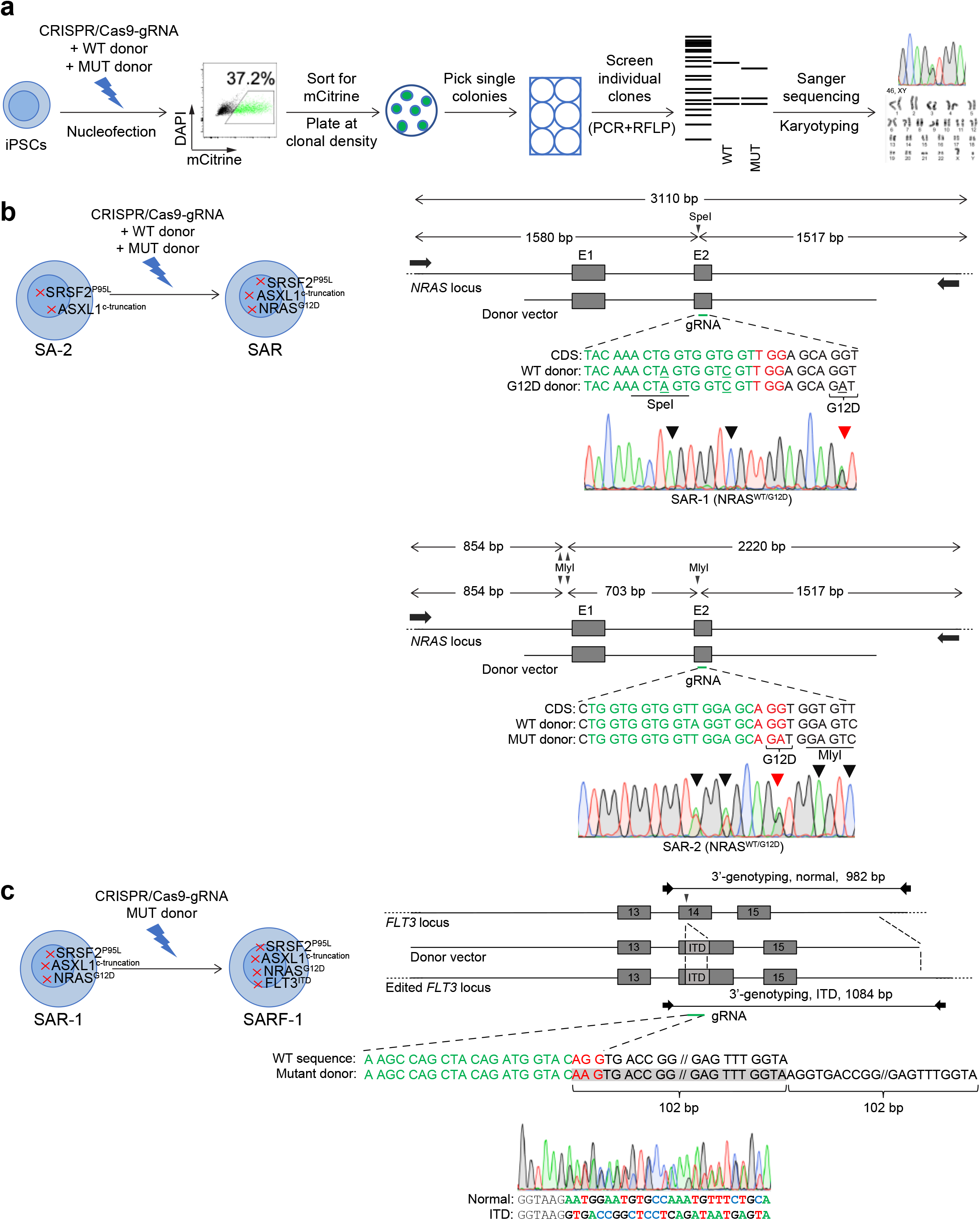
Generation of gene-edited iPSC lines. **a,** Scheme of the experimental strategy for generating CRISPR/Cas9-mediated gene edited clones. iPSCs were co-transfected with a plasmid expressing Cas9 (together with mCitrine) and the gRNA and donor DNA plasmids. Two donor plasmids (one carrying the hotspot mutation to be introduced and one the wild-type sequence) were co-delivered to facilitate selection of heterozygous mutant clones following selection of biallelically edited clones. mCitrine+ cells were sorted 48 hours after transfection by nucleofection to enrich for transfected and gene-edited cells and plated at clonal density. Individual clones were replated and initially screened by restriction fragment length polymorphism (RFLP) analysis and selected clones were further characterized by Sanger sequencing and karyotyping. Engineering of ASXL1 C-terminus truncations and of the SRSF2 P95L mutation have been previously described^12,13^. **b,** Strategy for introducing the *NRAS G12D* mutation in SA iPSCs. Upper panel from top to bottom: Schematic representation of the *NRAS* locus with the positions of the gRNA-1 target sequence, the SpeI restriction site and the PCR primers for RFLP analysis shown. A wild type (WT) donor template harbors two silent mutations (underlined), one to introduce a SpeI restriction site sequence for RFLP screening and one to prevent further cleavage by Cas9. A mutant (G12D) donor template harbors the same two silent mutations and additionally the *NRAS* c.35G>A mutation. The gRNA target sequence is shown in green and the PAM motif in red. Sanger sequencing of the *NRAS* locus in the SAR-1 line showing heterozygous *NRAS* c.35G>A mutation (red arrow) and the two silent mutations (black arrows) in homozygous state. (This sequencing result means that this line was generated through biallelic gene editing with one WT and one mutant donor). Lower panel from top to bottom: Schematic representation of the *NRAS* locus with the positions of the gRNA-2 target sequence, the MlyI restriction site and the PCR primers for RFLP analysis shown. A wild type (WT) donor template harbors four silent mutations (underlined), two to introduce a MlyI restriction site sequence for RFLP screening and two to prevent further cleavage by Cas9. A mutant (G12D) donor template harbors the same two silent mutations to introduce a MlyI restriction site sequence and additionally the *NRAS* c.35G>A mutation (that also inactivates further cleavage). The gRNA target sequence is shown in green and the PAM motif in red. Sanger sequencing of the *NRAS* locus in the SAR-2 line showing heterozygous *NRAS* c.35G>A mutation (red arrow) and the silent mutations (black arrows). **c,** Strategy for introducing the *FLT3-ITD* mutation in SAR iPSCs. From top to bottom: Scheme of the *FLT3* locus with the positions of the gRNA target sequence, the donor template introducing a 102 bp DNA sequence encoding the ITD and the primers used for PCR-based screening of edited colonies indicated. Donor template harboring the ITD and a silent mutation (underlined) to disrupt the PAM motif. The gRNA target sequence is shown in green and the PAM motif in red. Sanger sequencing of the edited SARF-3 line harboring monoallelic FLT3-ITD mutation.

**Extended Data Fig. 2.**
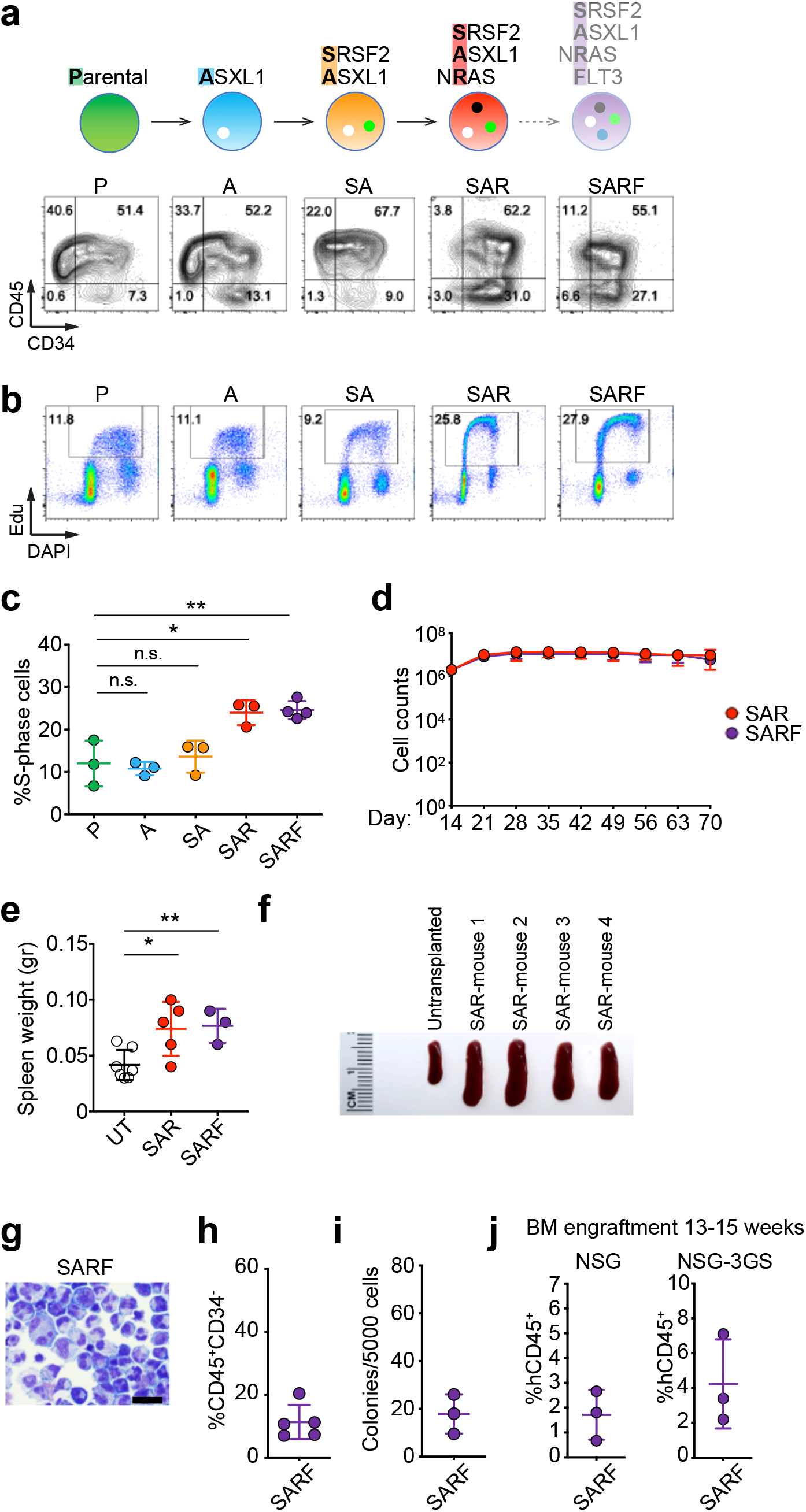
HSPCs differentiated from gene edited clonal iPSCs capture AML phenotypes in vitro and in vivo. **a,** Representative flow cytometry plots on day 14 of hematopoietic differentiation showing accumulation of CD34+ cells in SAR and SARF stages, consistent with a differentiation block. **b,** Representative cell-cycle analyses by flow cytometry of HSPCs differentiated from iPSC lines of the indicated genotypes. **c,** Cumulative cell-cycle analysis data of HSPC cells differentiated from one iPSC line for each stage (P: P-1, A: A-1, SA: SA-2, SAR: SAR-3, SARF: SARF-3). Mean and SEM from 3 independent differentiation experiments per line are shown. **d,** Cell counts of HSPCs at the indicated days of liquid hematopoietic differentiation culture. Mean and SEM of 2-3 independent differentiation experiments with 2 different iPSC lines from each genotype are shown. **e,** Spleen weight of untransplanted (UT) mice and mice transplanted with SAR or SARF cells 13 weeks posttransplantation. Mean and SEM of independent mice from one transplantation experiment are shown. **f,** Representative picture of spleens from 4 mice transplanted with SAR cells and one untransplanted mouse 13 weeks after transplantation. **g,** Wright-Giemsa staining of a representative cytospin preparation of hematopoietic cells derived from the SARF-3 iPSC line after 14 days of hematopoietic differentiation culture showing immature cells with blast morphology. Scale bars, 25 μm. **h,** Fraction of CD34^-^/CD45^+^ cells, i.e. hematopoietic cells that have lost CD34 expression upon maturation, on day 14 of hematopoietic differentiation. Mean and SEM of values from 5 independent differentiation experiments with 2 different SARF iPSC lines are shown. **i,** Number of colonies obtained from 5,000 cells seeded in methylcellulose assays on day 14 of hematopoietic differentiation. Mean and SEM of 3 independent methylcellulose experiments are shown. **j,** Levels of human engraftment in the BM of NSG and NSG-3GS mice 13 to15 weeks after transplantation with HSPCs derived from the SARF-3 iPSC line. Error bars show the mean and SEM of individual mice. (SARF: SRSF2^P95L^ /ASXL1^C-truncation^/ NRAS^G12D^/ FLT3^ITD^)

**Extended Data Fig. 3.**
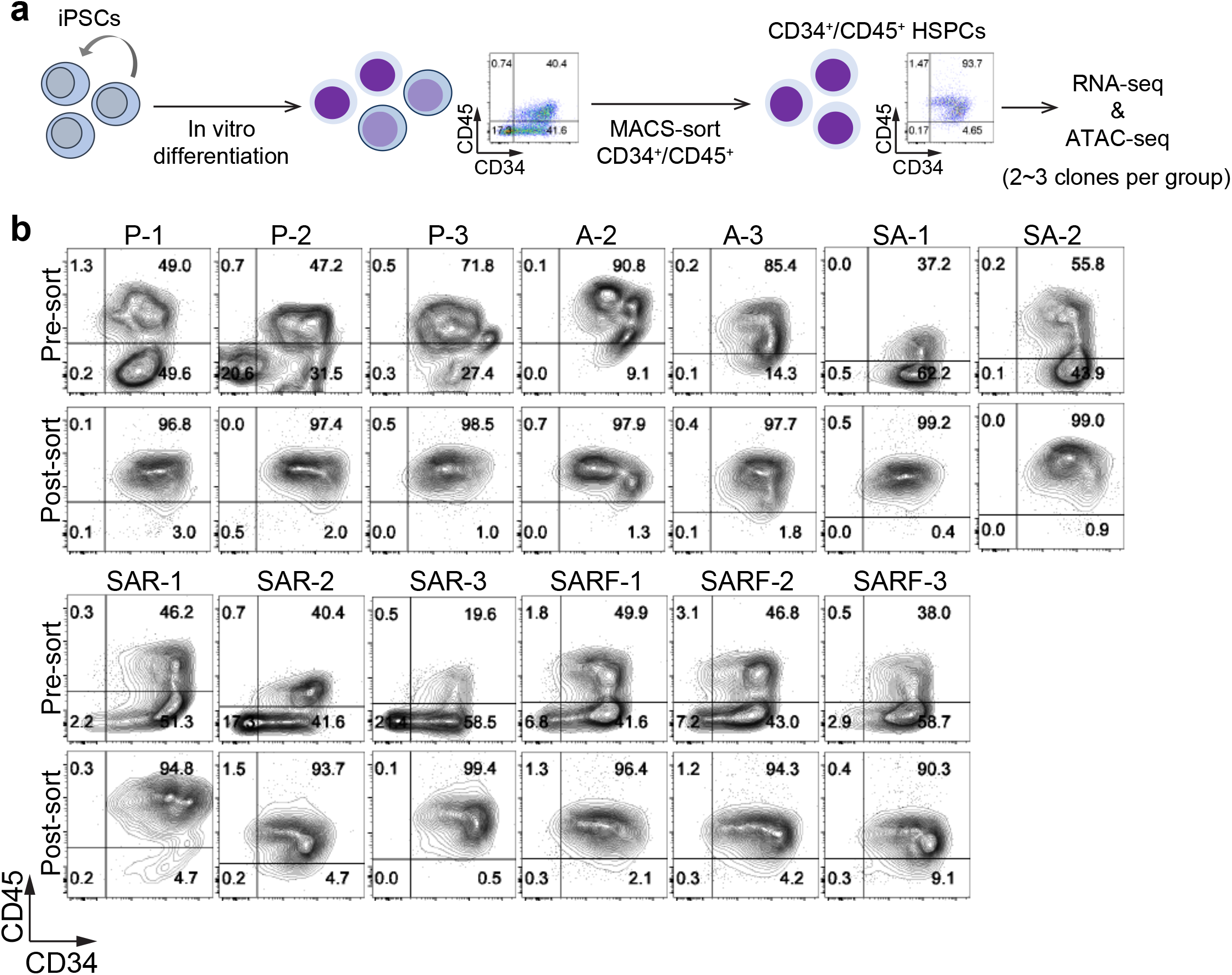
Isolation of CD34+/CD45+ iPSC-derived HSPCs for RNA- and ATAC-seq analyses. **a,** Scheme of isolation of iPSC-derived CD34+/CD45+ HSPCs for RNA- and ATAC-seq analyses using magnetic-activated cell sorting (MACS) with anti-CD45 beads on an empirically determined day of differentiation culture when nearly 100% of CD45+ cells are also still CD34+ (ranging from day 10 to 13, depending on the individual line and differentiation experiment). **b,** Flow cytometric assessment of cell purity of all MACS-sorted iPSC-HSPC samples used for the RNA-seq and ATAC-seq analyses.

**Extended Data Fig. 4.**
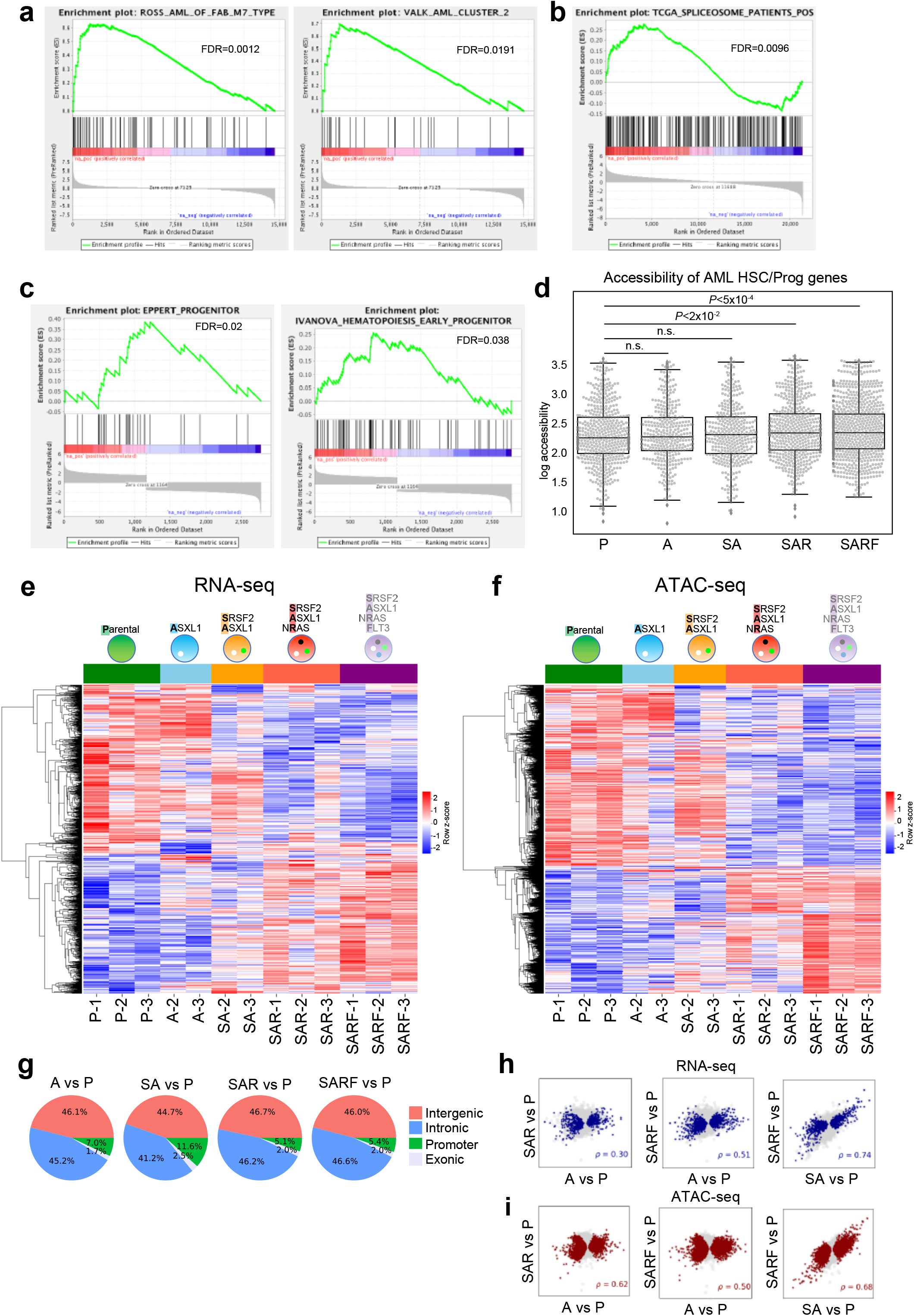
Gene expression and chromatin accessibility of gene-edited iPSC-HSPCs. **a,** GSEA plots of the indicated gene sets against the rank list of differentially expressed genes between SARF and P, showing enrichment of AML transcriptional programs in SARF cells (FDR<0.05). **b,** GSEA plot of a gene set derived from TCGA data from patients with chromatin and spliceosome gene mutations showing enrichment in genes that are associated with differentially accessible chromatin regions in SAR vs SA ranked by LFC accessibility change (FDR<0.05). **c,** GSEA plots showing enrichment of early hematopoietic progenitor gene sets in genes that are associated with differentially accessible chromatin regions in SARF vs P ranked by accessibility change (FDR<0.05). **d,** Beeswarm plot showing chromatin accessibility of all peaks associated with the 26 AML HSC/Prog genes from Fig. 2c in the indicated stages. The p values were calculated using one-sided unpaired Wilcoxon rank-sum tests. The average accessibility of the peaks of this gene set is significantly increased in SAR and SARF cells compared to P. **e,** Heatmap showing expression of all genes that were differentially expressed in at least one of the comparisons A vs P, SA vs P, SAR vs P, SARF vs P (log2FC > 1, FDR-adjusted p < 0.05). **f,** Heatmap showing accessibility of all peaks that were differentially accessible in at least one of the comparisons A vs P, SA vs P, SAR vs P, SARF vs P (FDR-adjusted p < 0.05). **g,** Genomic distribution of differentially accessible peaks in the indicated comparisons. **h,** Scatter plots of gene expression changes (log2FC) showing correlation between the two comparisons indicated in the x and y axes of each plot. Significant changes (FDR-adjusted p < 0.05) in the x-axis comparisons are highlighted in blue. The ρ values represent Spearman correlation values calculated from the log2FC of genes with significant changes in the x-axis comparison. **i,** Scatter plots of chromatin accessibility changes (log2FC) showing correlation between the two comparisons indicated in the x and y axes of each plot. Peaks with a significant change (FDR-adjusted p < 0.05) in the x-axis comparisons are highlighted in red. The ρ values are Spearman correlation values calculated from the log2FC of peaks with significant changes in the x-axis comparison.

**Extended Data Fig. 5.**
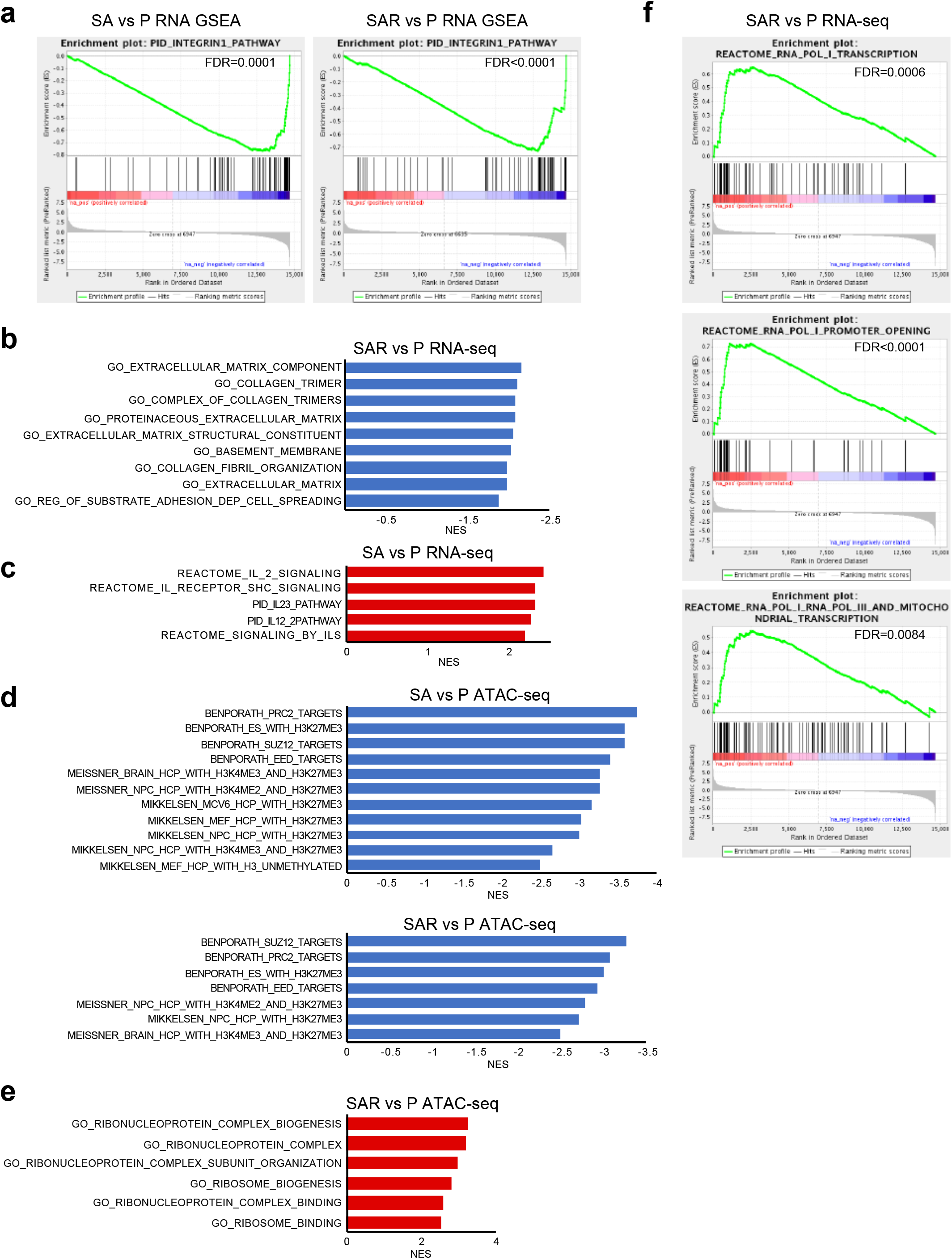
GO terms and gene sets enriched in distinct stages. **a,** GSEA plots of the indicated gene sets against the rank list of differentially expressed genes in SA vs P and SAR vs P, showing depletion of genes implicated in integrin pathway in SA and SAR cells (FDR<0.05). **b,** Gene-ontology (GO) terms related to adhesion and extracellular matrix (ECM) enriched in genes downregulated in SAR vs P (NES: Normalized enrichment score, FDR<0.05). **c,** GSEA results showing enrichment of interleukin signaling gene sets in genes upregulated in SA vs P (NES: Normalized enrichment score, FDR<0.05). **d,** GSEA results showing enrichment of histone methylation-related gene sets in genes downregulated in SA vs P and SAR vs P (NES: Normalized enrichment score, FDR<0.05). **e,** GO terms related to ribosome biogenesis enriched in genes associated with peaks more accessible in SAR vs P cells (NES: Normalized enrichment score, FDR<0.05). **f,** GSEA plots of the indicated gene sets against the rank list of differentially expressed genes between SAR and P, showing enrichment of genes implicated in RNA PolI transcription in SAR cells (FDR<0.05).

**Extended Data Fig. 6.**
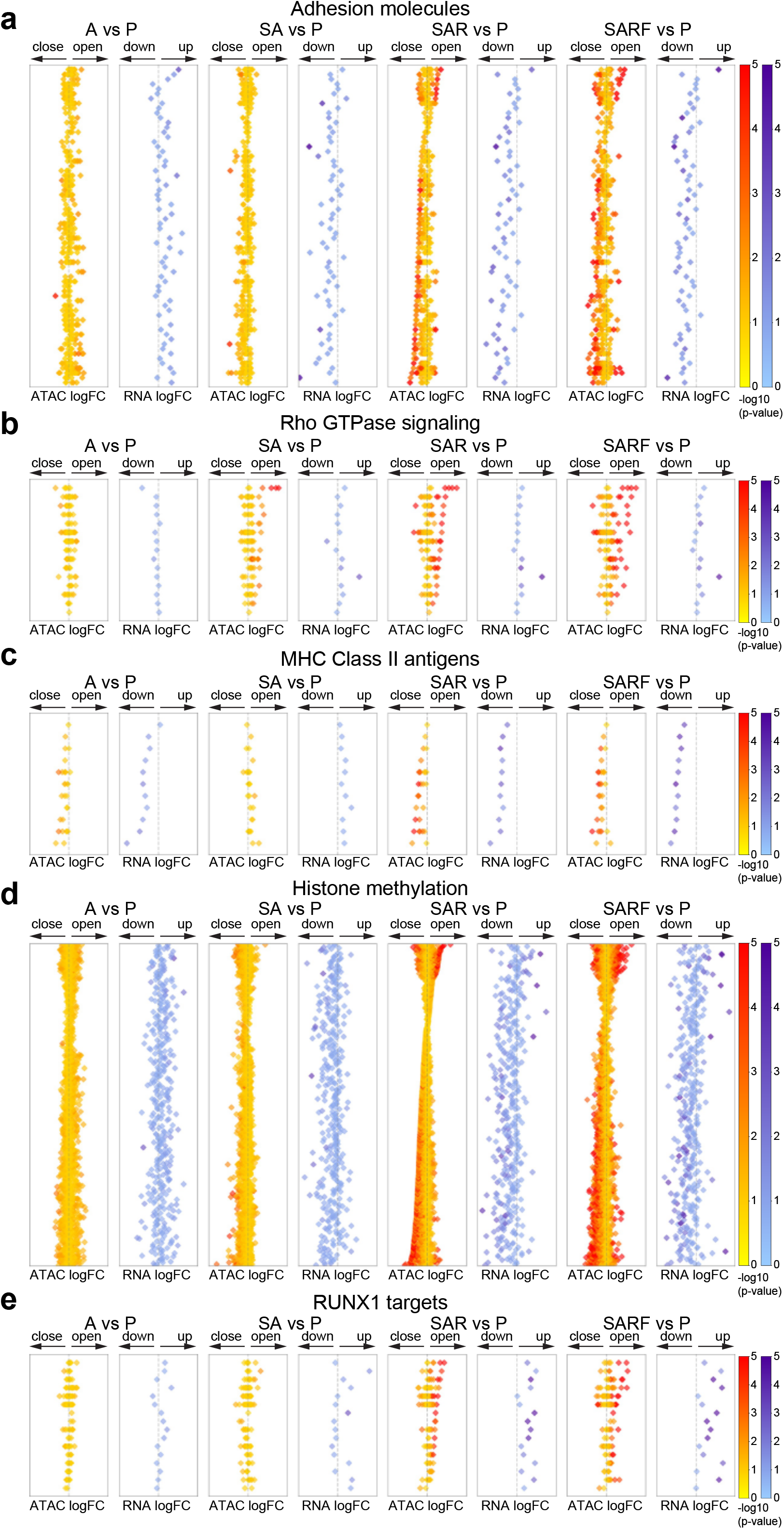
Diamond plots of gene expression and chromatin accessibility changes of specific gene sets across. **a-e**, Diamond plots showing the expression change (log2FC) of genes from the indicated gene sets together with the accessibility change (log2FC) of the peaks associated with them in the indicated comparisons (A vs P, SA vs P, SAR vs P and SARF vs P). Each row represents one gene and all peaks associated with it.

**Extended Data Fig. 7.**
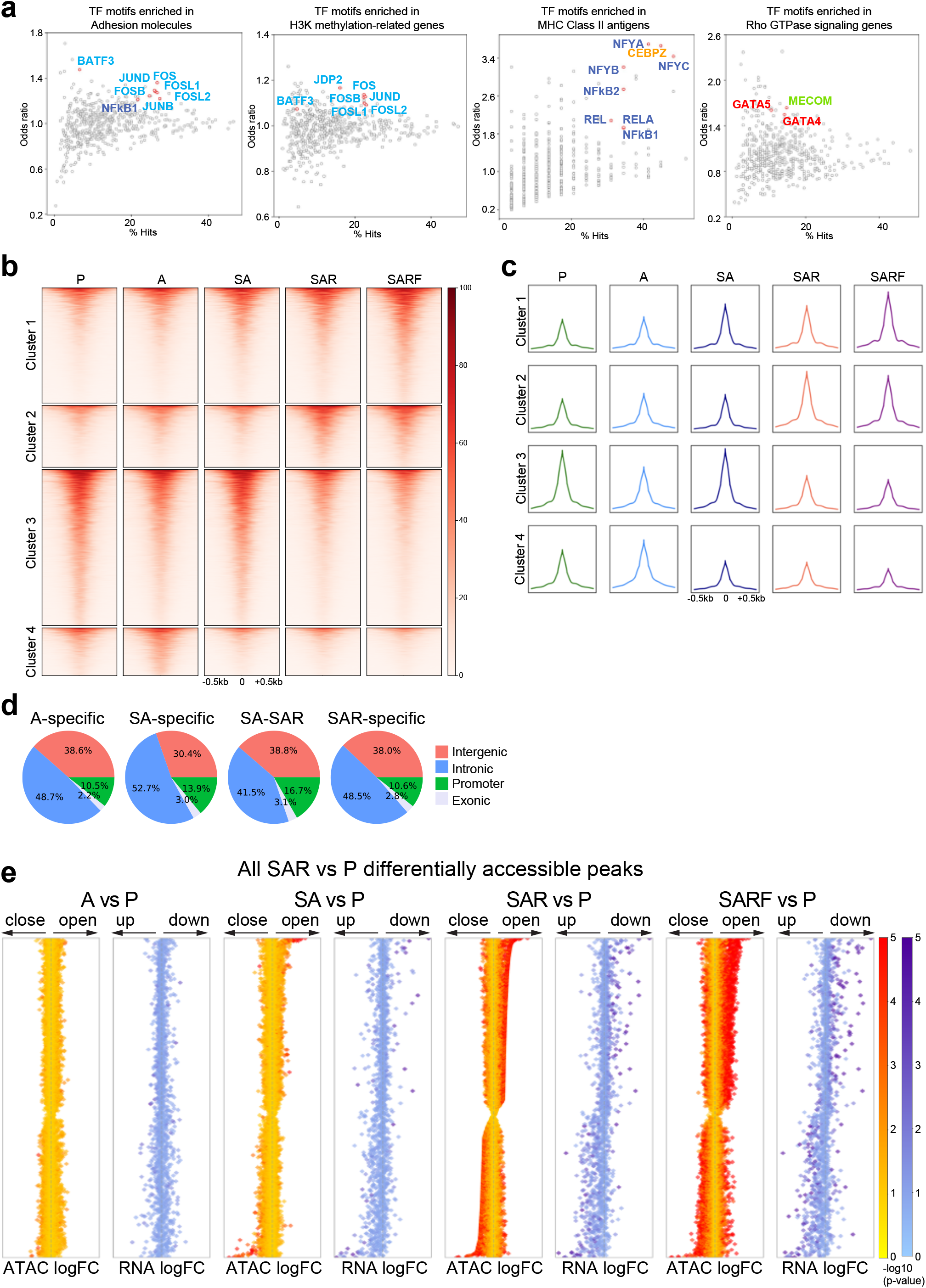
Stage-specific chromatin accessibility analyses. **a,** TF motifs enriched in the indicated gene sets. The x axis indicates the percentage of hits for each TF motif in peaks associated with each annotated gene set. The y axis indicates motif enrichment for the peaks associated with each gene set compared to background (total atlas, which comprises all reproducible ATAC-seq peaks from all stages), measured as odds ratio, with selected TFs with enriched motifs highlighted in red. AP-1 family (FOS, FOSB, FOSL1, FOSL2, JUNB, JUND, BATF3) and NFkB1 motifs are enriched in genes encoding adhesion molecules. Histone lysine methylation-related genes show enrichment for AP-1 motifs (FOS, FOSB, FOSL1, FOSL2, JUND, BATF3, JDP2). MHC Class II genes show enrichment for NFkB (NFYA, NFYB, NFYC, NFkB1, NFkB2, REL, RELA) and CEBP motifs. GATA and MECOM TF motifs are enriched in genes associated with Rho GTPase signaling. **b,** Tornado plot showing all peaks that are differentially accessible (p <0.05) in at least one of the comparisons A vs P, SA vs P, SAR vs P, SARF vs P, with four clusters defined by hierarchical clustering shown. **c,** Average of normalized counts from the tornado plot shown in b, visualized as metapeaks. **d,** Genomic distribution of stage-specific peaks from Fig. 4c with the percentage of intergenic, intronic, promoter and exonic genomic regions shown. **e,** Diamond plots showing the accessibility change (log2FC) of the all peaks that are differentially accessible between SAR and P together with the expression change (log2FC) of genes associated with them. Each row represents one gene and all peaks associated with it across all comparisons (A vs P, SA vs P, SAR vs P and SARF vs P), allowing back- and forward-tracking of changes in successive stages.

**Supplementary Table 1.** List of all isogenic iPSC lines used in this study.

| Line | Genotype |  | Parental line | gRNA | Reference |
| --- | --- | --- | --- | --- | --- |
| P-1 |  | WT (N-2.12-D-1-1) | N/A | N/A | Kotini et al., 2015 |
| P-2 |  | WT (N-2.12, clone WT-U83) | N/A | N/A | Unpublished |
| P-3 |  | WT (N-2.12-GFP) | N/A | N/A | Kotini et al., 2015 |
| A-1 |  | ASXL1(WT/c.1888_1910del23) | P-1 | ASXL1-gRNA-2 | Kotini et al., 2017 |
| A-2 |  | ASXL1(WT/c.1888_1910del23) | P-1 | ASXL1-gRNA-2 | Kotini et al., 2017 |
| A-3 |  | ASXL1(WT/c.1900_1901del2) | P-1 | ASXL1-gRNA-1 | Kotini et al., 2017 |
| SA-1 |  | SRSF2(WT/c.284C>T), ASXL1(WT/c.1901_1902insA) | S-1 | ASXL1-gRNA-2 | This study |
| SA-2 |  | SRSF2(WT/c.284C>T), ASXL1(WT/c.1888_1910del23) | S-1 | ASXL1-gRNA-2 | This study |
| SA-3 |  | SRSF2(WT/c.284C>T), ASXL1(WT/c.1888_1910del23) | S-1 | ASXL1-gRNA-2 | This study |
| SAR-1 |  | SRSF2(WT/c.284C>T), ASXL1(WT/c.1888_1910del23), NRAS(WT/c.35G>A) | SA-2 | NRAS-gRNA-1 | This study |
| SAR-2 |  | SRSF2(WT/c.284C>T), ASXL1(WT/c.1888_1910del23), NRAS(WT/c.35G>A) | SA-2 | NRAS-gRNA-2 | This study |
| SAR-3 |  | SRSF2(WT/c.284C>T), ASXL1(WT/c.1888_1910del23), NRAS(WT/c.35G>A) | SA-2 | NRAS-gRNA-2 | This study |
| SARF-1 |  | SRSF2(WT/c.284C>T), ASXL1(WT/c.1888_1910del23), NRAS(WT/c.35G>A), FLT3(WT/ITD) | SAR-1 | FLT3-gRNA-1 | This study |
| SARF-2 |  | SRSF2(WT/c.284C>T), ASXL1(WT/c.1888_1910del23), NRAS(WT/c.35G>A), FLT3(WT/ITD) | SAR-1 | FLT3-gRNA-1 | This study |
| SARF-3 |  | SRSF2(WT/c.284C>T), ASXL1(WT/c.1888_1910del23), NRAS(WT/c.35G>A), FLT3(WT/ITD) | SAR-1 | FLT3-gRNA-1 | This study |

**Supplementary Table 2.**
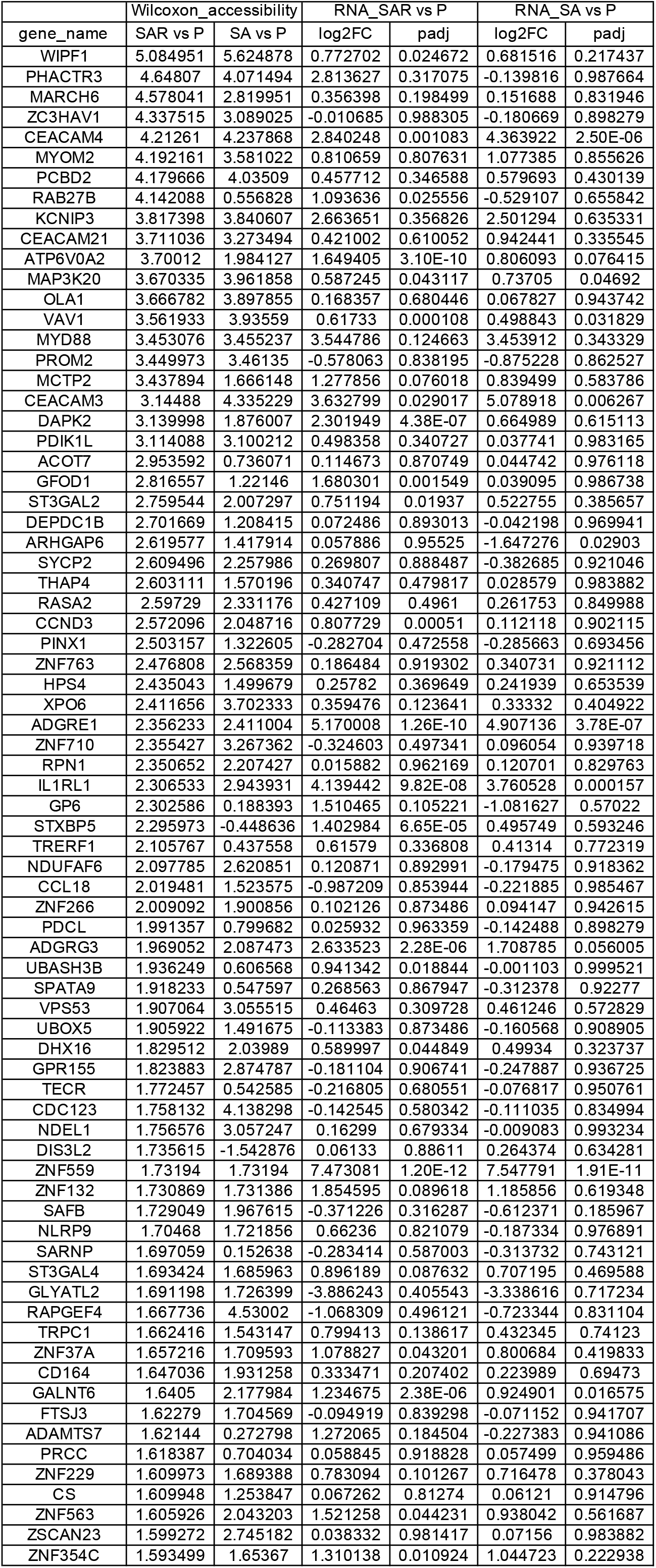
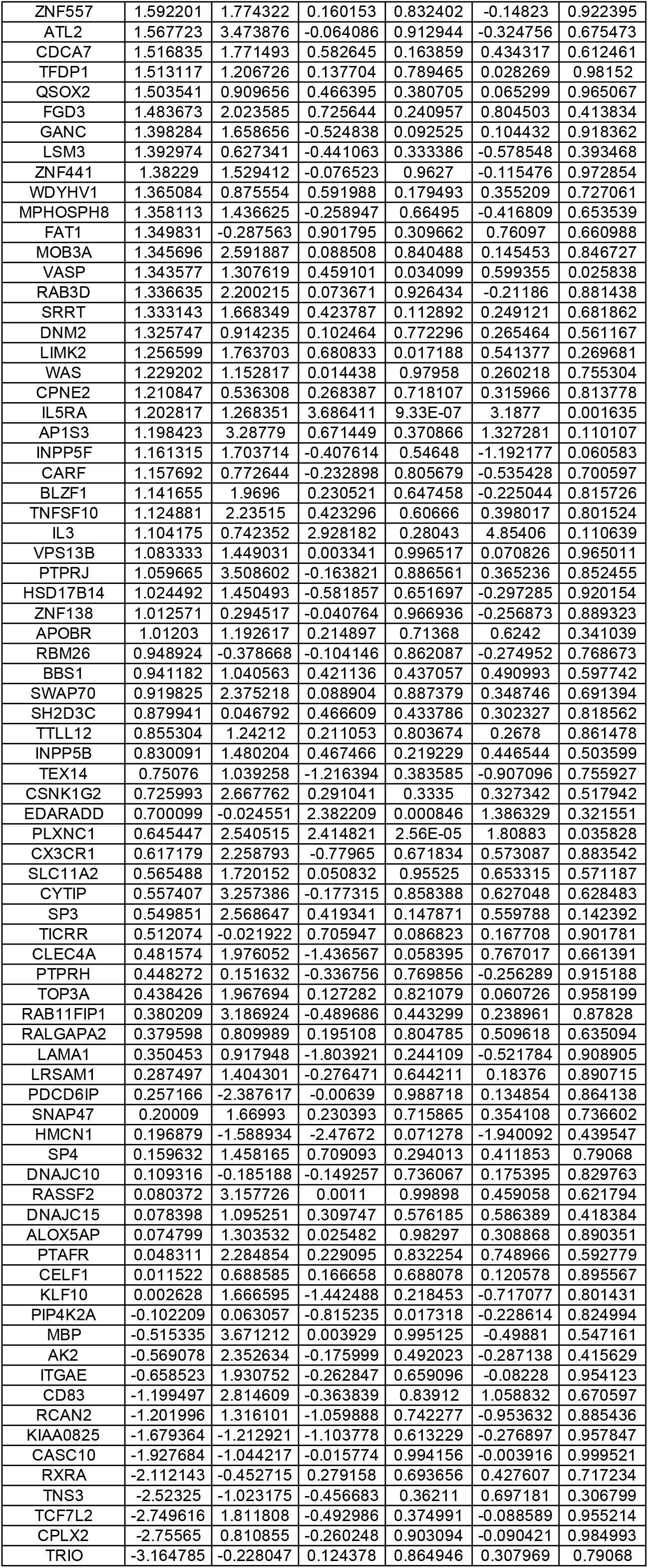
List of all genes associated with peaks of the SA-SAR group.

**Supplementary Table 3.** List of “AML-early genes”.

| gene_name | Wilcoxon_accessibility |  | RNA_SAR vs P |  | RNA_SA vs P |  |
| --- | --- | --- | --- | --- | --- | --- |
|  | SAR vs P | SA vs P | log2FC | padj | log2FC | padj |
| CEACAM4 | 4.21261039 | 4.23786811 | 2.84024803 | 0.00108254 | 4.36392227 | 2.50E-06 |
| ATP6V0A2 | 3.7001202 | 1.98412708 | 1.64940528 | 3.10E-10 | 0.80609312 | 0.07641492 |
| MAP3K20 | 3.67033537 | 3.96185838 | 0.5872446 | 0.04311676 | 0.73705028 | 0.04691965 |
| VAV1 | 3.56193258 | 3.93558989 | 0.61733005 | 0.0001082 | 0.49884259 | 0.03182889 |
| CEACAM3 | 3.14487984 | 4.33522905 | 3.63279862 | 0.02901655 | 5.07891772 | 0.00626675 |
| ADGRE1 | 2.35623343 | 2.41100405 | 5.17000811 | 1.26E-10 | 4.90713591 | 3.78E-07 |
| IL1RL1 | 2.30653281 | 2.9439312 | 4.13944207 | 9.82E-08 | 3.76052786 | 0.00015735 |
| ADGRG3 | 1.96905194 | 2.08747348 | 2.6335228 | 2.28E-06 | 1.7087847 | 0.05600491 |
| ZNF559 | 1.73194 | 1.73194 | 7.4730813 | 1.20E-12 | 7.54779146 | 1.91E-11 |
| GALNT6 | 1.6405001 | 2.17798434 | 1.23467454 | 2.38E-06 | 0.92490084 | 0.01657481 |
| VASP | 1.34357699 | 1.30761899 | 0.45910147 | 0.03409906 | 0.59935463 | 0.0258383 |
| IL5RA | 1.20281711 | 1.26835068 | 3.68641087 | 9.33E-07 | 3.18770018 | 0.00163531 |
| PLXNC1 | 0.64544677 | 2.54051479 | 2.41482101 | 2.56E-05 | 1.80882968 | 0.0358282 |

## Methods

### CRISPR-Cas9 gene editing of human iPSCs

We used the previously described normal iPSC line N-2.12-D-1-1 as the parental line^11^. We used CRISPR/Cas9-mediated non-homology end-joining (NEJM) to introduce the ASXL1 C-terminus truncation and homology-directed repair (HDR) for all other mutations. Engineering of ASXL1 C-terminus truncations and of SRSF2 P95L mutation were performed as previously described^12,13^. Multiple independent clones with the desired mutation were isolated after each gene editing step and, following genetic and preliminary phenotypic characterization to exclude potential outliers, one clone was selected for the subsequent step (Supplementary Table 1).

To introduce the NRAS G12D mutation or the FLT3-ITD mutation, we used a plasmid expressing gRNAs under the U6 promoter and Cas9 linked to mCitrine with a P2A driven by the CMV promoter^13^ and donor DNA plasmids. For NRAS G12D, two different gRNAs targeting the NRAS locus within the second exon (cutting site between 5 bp and 12 bp from the 35 G>A mutation site, sequences shown in Extended Data Fig. 1b) were designed, assembled by a two-step overlapping PCR reaction downstream of the U6 promoter sequence and cloned in the gRNA/Cas9 plasmid. Two sets (one for each gRNA) of two donor DNA plasmids, one containing the G12D mutation and one the corresponding wild-type (WT) sequence, containing 5’ (1040 bp) and 3’ (1040 bp) homology arms consisting of nucleotides 115258748 – 115259787 and 115257707– 115258746 (hg19 human genome assembly), respectively, were constructed. The donor plasmids also contained silent mutations (shown in Extended Data Fig. 1b) to introduce a new restriction site sequence (SpeI or MlyI, respectively) and to prevent further cleavage by Cas9. The entire 5’+3’ homology sequence was amplified from N-2.12 genomic DNA and the 35 G>A and/or silent mutations to introduce new restriction enzyme recognition sites and to prevent cleavage by Cas9 were introduced by two-step overlapping PCR before subsequent cloning into the donor plasmid.

To generate SRSF2^P95L^/ASXL1^c-truncation^/NRAS^G12D^ (SAR) iPSCs, the SA-2 iPSC line was cultured in hESC media containing 10 mM Y-27632 for at least one hour before nucleofection. The cells were dissociated into single cells with accutase and 1 million cells were used for nucleofection with 5 μg of gRNA/Cas9 plasmid and 5 μg of each donor plasmid (WT and G12D) using Nucleofector II (Lonza) and program B-16. Immediately after nucleofection the cells were replated on MEFs. mCitrine+ cells were FACS-sorted 48 hr after transfection and plated as single cells at clonal density (1000 FACS-sorted cells per 60-mm dish). After 10-12 days, single colonies were picked in separate wells of a 6-well plate, allowed to grow for approximately 3-6 days and screened by PCR. 1-3 medium-sized colonies from each individual clone were picked directly into a 0.2 ml tube, pelleted and lysed. Restriction Fragment Length Polymorphism (RFLP) analysis was performed after PCR with primers F: TCTGAGGACCATATGAGGGTAGA and R: TGGTTTTATGGCAACAGGACT and digestion of the product with MlyI or SpeI. Bi-allelically targeted clones were selected and the PCR products were cloned into the PCR-4 TOPO TA vector (Invitrogen) and sequenced to select clones heterozygous the NRAS^G12D^ mutation.

To generate SRSF2^P95L^/ASXL1^c-truncation^/NRAS^G12D^/FLT3^ITD^ (SARF) iPSCs, a 102 bp sequence containing the FLT3-ITD mutation (duplication of 102 bp encoding part of the juxtamembrane domain) was inserted in the SAR-1 line as follows. A gRNA targeting the FLT3 locus within the 14^th^ exon (cutting site 3 bp from the duplication insertion site, sequences shown in Extended Data Fig. 1c) was designed and cloned as above. A donor template containing a 5’ homology arm (923 bp), the ITD sequence (102 bp) and a 3’ homology arm (714 bp), consisting, respectively, of nucleotides 28608318 – 28609240, 28608216 – 28608317 and 28607604 – 28608317 (hg19 human genome assembly), was assembled and cloned in a plasmid. The SAR-1 iPSC line was nucleofected with 5 μg of gRNA/Cas9 plasmid and 10 μg of the donor plasmid PCR as described above and screened by PCR with primers F: CACTCTTTTGTTGCAGGCCC and R: CGGCAACCTGGATTGAGACT. Mono-allelically targeted clones were selected and the PCR products were cloned into the PCR-4 TOPO TA vector (Invitrogen) and sequenced. Clones with one *FLT3^ITD^* allele and one wild-type intact (without indels) allele were selected. To ensure clonality, an additional step of single-cell cloning was performed after each gene editing step.

### Human iPSC culture, hematopoietic differentiation and in vitro phenotypic characterization

Culture of human iPSCs on mitotically inactivated MEFs or feeder-free conditions, was performed as previously described^13^. Hematopoietic differentiation was performed using a spin-EB protocol previously described^13^. In the end of the differentiation culture, the cells were collected and dissociated with accutase into single cells and used for flow cytometry, cytological analyses or clonogenic assays, as described^13^. Competitive growth assays were performed using an isogenic GFP-marked iPSC line (N-2.12-GFP), as previously described^13^. For cell cycle analyses, 500,000 cells were incubated with EdU added to the media for 4 hours and analyzed with the Click-iT™ EdU Alexa Fluor™ 488 Flow Cytometry Assay Kit.

### Flow cytometry and cell sorting

The following antibodies were used: CD34-PE (clone 563, BD Pharmingen), CD45-APC (clone HI30, BD PharMingen), CD33-BV421 (clone WM53, BD Horizon), mCD45-PE-Cy7 (clone 30-F11, BD Biosciences), CD19-PE (clone HIB19, BD Pharmingen). Cell viability was assessed with DAPI (Life Technologies). Cells were assayed on a BD Fortessa and data were analyzed with FlowJo software (Tree Star). Cell sorting was performed on a BD FACS Aria II.

### Transplantation into NSG mice

NSG (NOD.*Cg-Prkdc^scid^Il2rg^tm1Wjl^*/SzJ) and NSG-SGM3 (NOD.*Cg-Prkdc^scid^Il2rg^tm1Wjl^*Tg(CMV-IL3,CSF2,KITLG)1Eav/MloySzJ) mice were purchased from Jackson Laboratories and housed at the Center for Comparative Medicine and Surgery at Icahn School of Medicine at Mount Sinai. One day before transplantation, the mice were injected intraperitoneally with 30mg/kg Busulfan solution. P, A, SA, SAR and SARF iPSC-HSPCs from day 13-15 of hematopoietic differentiation were resuspended in StemPro-34 and injected via the tail vein using a 25G needle at 1×10^6^ cells per 100μL per mouse. The mice were sacrificed after 13-15 weeks. Bone marrow was collected from the femurs and tibia for flow cytometry and spleens were harvested and their weight recorded. All mouse studies were performed in compliance with Icahn School of Medicine at Mount Sinai laboratory animal care regulations.

### RNA sequencing

Two to three clones of P, A, SA, SAR and SARF iPSCs were subjected to hematopoietic differentiation. Magnetic cell sorting of CD45^+^ cells was performed using the MACS cell separation microbeads and reagents (Miltenyi Biotec). Total RNA was extracted with the RNeasy mini kit (Qiagen). PolyA-tailed mRNA was selected with beads from 1μg total RNA using the NEBNext Poly(A) mRNA Magnetic Isolation Module (New England Biolabs). cDNAs were generated using random hexamers and ligated to barcoded Illumina adaptors with the NEXTflex Rapid Directional RNA-Seq Library Prep Kit (Bioo Scientific). Sequencing of 75 nucleotide-long single-end reads was performed in a NextSeq-500 (Illumina).

### ATAC sequencing

50,000 MACS-sorted CD45^+^ cells from two to three clones of P, A, SA, SAR and SARF cells were processed as follows: nuclei were isolated by lysing with 50 ul of ATAC lysis buffer (10 mM Tris pH 7.4, 10 mM NaCl, 3 mM MgCl2, 0.1% NP40, 0.1% Tween-20, and 0.01% Digitonin) and washing with 1 ml of ATAC wash buffer (10 mM Tris pH 7.4, 10 mM NaCl, 3 mM MgCl2, 0.1% Tween-20). Cell lysates were spun to obtain nuclear pellets, which were subjected to transposase reaction using the Illumina Nextera DNA Sample Preparation Kit according to the manufacturer’s instructions. The final libraries were quantified using the Agilent BioAnalyzer. Sequencing of 75 nucleotide-long paired-end reads was performed in a NextSeq-500 (Illumina).

### Differential gene expression analysis of RNA-seq data

Fastq files were aligned to GRCh37.75 (hg19) reference genome and RefSeq gene annotation using STAR alignment tool. Aligned bam files were filtered for uniquely aligned reads with MAPQ > 10. Subsequently, RNA-seq reads were counted using GRCh37.75 annotation with tRNA and rRNA removed. Read counting in exonic regions was computed using SummarizeOverlaps function from R-package “GenomicAlignments” with IntersectionNotEmpty mode. There were 10-20 million reads for each sample. We filtered genes with FPKM < 0.1 in at least 2 samples and count < 10 in at least 2 samples. There were 15,115 genes in the count table after filtering. Differential expression analysis was performed using R-package DESeq2. Genes with an FDR cutoff of 0.05 were considered significantly differentially expressed. Subsequent principal component and heatmap analyses were computed with genes that were significantly differentially expressed between conditions A, SA, SAR, or SARF vs P. Normalized counts were log2-transformed and row z-score normalized prior to heatmap generation. To generate the summary gene expression dynamics shown in Fig. 3m, the median normalized counts per gene set were used prior to log transformation.

### Differential accessibility analysis of ATAC-seq data

Fastq files were trimmed with Trimmomatic to remove adapter sequences and aligned to GRCh37.75 (hg19) reference genome using Bowtie2 alignment tool. Subsequently, low quality PE reads with MAPQ < 33 were removed using samtools and duplicate reads were removed using Picard. All aligned reads were shifted to removeTn5 transposase artifacts, as described^35^. We then called peaks for each replicate using MACS2 with the following parameters (-p 1e-2 --nomodel --shift 0 --extsize 76). For each pair of replicates, we filtered peaks using an Irreproducible Discovery Rate (IDR) cutoff of 0.05. Finally, we created an ATAC-seq atlas of 93,782 peaks by combining all reproducible peaks using the bedtools merge tool. Peaks were annotated to the nearest gene using the RefSeq database.

Differential accessibility analysis was performed using R-package DESeq2. As in the RNA-seq analysis, peaks with an FDR cutoff of 0.05 were considered significantly differentially accessible, and subsequent PCA and heatmap analyses were computed with peaks significantly differentially accessible between conditions A, SA, SAR, or SARF vs P. We considered a gene to be significantly differentially accessible in a transition if the log fold changes of the peaks annotated to that gene were significantly higher as compared to the log fold changes of the background atlas of peaks, as determined by a Wilcoxon rank sum test. Swarm plots were generated using these gene-based accessibility values for a list of AML HSC/Prog genes at each condition using the swarmplot function in the python seaborn package (https://seaborn.pydata.org).

### GSEA and GO analyses

GSEA analyses was performed on all 4,733 curated gene sets and GO analyses was performed on all 4,079 GO gene sets in the Molecular Signatures Database (MSigDB, http://www.broadinstitute.org/msigdb). Significantly enriched gene sets were defined by cutoff FDR<0.05.

Additionally, we obtained unprocessed RNA-seq read counts from NCI’s Genomic Data Commons Data Portal (TCGA-LAML) and grouped patients as “chromatin-spliceosome subgroup” if they were found to have *SRSF2* mutations^36^ coexisting with mutations in chromatin regulators (n=15). We performed DESeq2 differential expression analysis and defined a set of genes significantly upregulated in this subgroup (“chromatin-spliceosome” gene set) (LFC > 0, FDR-adjusted p < 0.05).

### Motif enrichment analysis

We used FIMO to search for motif hits in the ATAC-seq atlas with default parameters using CISBP Single Species DNA Homo_sapiens motif database (http://meme-suite.org/db/motifs), restricted to expressed TFs only, based on our RNA-seq filtering thresholds. For each motif, we compared accessibility log2FC distribution of peaks containing the motif with the accessibility log2FC of the atlas by KS test, as described previously^37^. The KS statistic was then plotted against the percentage of motif hits in the atlas. Positive/negative sign of KS statistic indicates the direction of change with positive indicating increased accessibility and negative indicating decreased accessibility. Motif enrichment odds ratio plots were generated for a given gene set by plotting the percentage of motif hits in peaks annotated to that gene set on the x-axis and the odds ratio of a motif being annotated to that gene set on the y-axis. De novo motif analysis was done using HOMER findMotifsGenome.pl where a set of peaks of interest was compared to the background atlas of peaks.

### Correlation of chromatin accessibility to normal hematopoiesis

To compare ATAC-seq data to normal hematopoiesis, we obtained bulk ATAC-seq data from Corces et al.^18^, including 590,650 peaks. Of the 88,748 peaks that overlapped between the two datasets, we took peaks that were significantly differentially accessible and had a log fold change cutoff of 1 between SAR and P. We then computed pairwise Pearson correlations between the normalized reads in P, A, SA, SAR, SARF and the normal hematopoietic progenitor populations HSC, MPP, CMP and GMP.

### Definition and visualization of ATAC-seq stage-specific groups

To define stage-specific groups of peaks in ATAC-seq as shown in Fig. 4c, we implemented a generalized linear model using DESeq2 to identify peaks following specified up/down patterns across stages. In this way, we found peaks whose accessibility in any stage belonging to the ‘‘up’’ set showed greater magnitude than any of those in the ‘‘down’’ set, with an FDR cutoff of 0.1. For example, the SA specific group had only SA as up and P, A, and SAR as down, whereas the SA-SAR group had SA and SAR as up and P and A as down. Tornado plots were generated using the bigWig file format for each stage, averaged across replicates. The 99^th^ percentile of the bigWig file readout of the ATAC-seq data was used to set the maximum color in the tornado plots to reduce the outlier effect of highly accessible peaks, as described^37^. The peaks were centered on their summit and sorted by their mean signal. Meta-peak plots were generated by aggregating the signal along the y-axis of the tornado plots. Motif enrichment in stage-specific groups was calculated with a hypergeometric test where the proportion of peaks containing a transcription factor motif as identified by FIMO was compared to the background atlas.

### Statistical analysis

Statistical analysis was performed with GraphPad Prism software. Pairwise comparisons between different groups were performed using a two-sided unpaired unequal variance t-test.

For all analyses, p < 0.05 was considered statistically significant. Investigators were not blinded to the different groups.

### iPSC generation from an AML patient

A BM sample from an AML patient was obtained with informed consent under a protocol approved by a local Institutional Review Board. Cryopreserved BM mononuclear cells were thawed and cultured in X-VIVO 15 media with 1% nonessential amino acids (NEAA), 1 mM L-glutamine and 0.1 mM β-mercaptoethanol (2ME) and supplemented with 100 ng/ml stem cell factor (SCF), 100 ng/ml Flt3 ligand (Flt3L), 100 ng/ml thrombopoietin (TPO) and 20 ng/ml IL-3 for 1-4 days. An aliquot of the cells before culture was used for gene panel sequencing with a custom capture bait set including the coding regions of 163 myeloid malignancy genes and 1,118 genome-wide single nucleotide polymorphism (SNP) probes for copy number analysis, with on average one SNP probe every 3 Mb. Bait tiling was conducted at 2x.

For induction of reprogramming, 200,000 cells were transduced with the viral cocktail CytoTune-iPS 2.0 Sendai reprogramming kit (Invitrogen), containing *KLF4, OCT4,* and *SOX2* (KOS) virus, the *c-MYC* virus and the *KLF4* virus at an MOI ratio of 5:5:3 *(KOS: MYC: KLF4),* in a total volume of 400 μL. 24 hours after transduction 1 mL of X-VIVO 15 with cytokines was added. 24 hours later, the cells were harvested and plated on mitotically inactivated MEFs in 6-well plates and the plates were centrifuged at 500g for 30 min at RT. The next day and every day thereof, half of the medium was changed to hESC medium with 0.5 mM valproic acid (VPA). Colonies with hPSC morphology were manually picked and expanded, as described.

### VAF analysis

Cells treated with AraC, bafilomycin A1 or DMSO were collected before and after 48 hours of treatment. Genomic DNA was isolated and the *SRSF2* and *CSF3R* genomic loci were amplified in 20 PCR cycles. The universal NEXTflex adapters were tagged in a second round of 10 cycles of PCR amplification and the amplicons were purified and sequenced in the Illumina platform. Sequencing reads (75 nt) were sorted into mutant and wild type amplicons and the VAF was calculated as the ratio of mutant amplicon reads versus the sum of mutant and wild type amplicon reads.

### Data availability

RNA-sequencing and ATAC-sequencing data of this study have been deposited in GEO with the accession code TBD. Files can be accessed using this link: https://dataview.ncbi.nlm.nih.gov/object/PRJNA594644?reviewer=ra329es32nnnbo984edlpjjlim

### Code availability

N/A

